# MS Binding Assays with UNC0642 as reporter ligand for the MB327 binding site of the nicotinic acetylcholine receptor

**DOI:** 10.1101/2023.11.15.567260

**Authors:** Valentin Nitsche, Georg Höfner, Jesko Kaiser, Christoph G.W. Gertzen, Thomas Seeger, Karin V. Niessen, Dirk Steinritz, Franz Worek, Holger Gohlke, Franz F. Paintner, Klaus T. Wanner

## Abstract

Intoxications with organophosphorus compounds (OPCs) based chemical warfare agents and insecticides may result in a detrimental overstimulation of muscarinic and nicotinic acetylcholine receptors evolving into a cholinergic crisis leading to death due to respiratory failure. In the case of the nicotinic acetylcholine receptor (nAChR), overstimulation leads to a desensitization of the receptor, which cannot be pharmacologically treated so far. Still, compounds interacting with the MB327 binding site of the nAChR like the bispyridinium salt MB327 have been found to re-establish the functional activity of the desensitized receptor. Only recently, a series of quinazoline derivatives with UNC0642 as one of the most prominent representatives has been identified to address the MB327 binding site of the nAChR as well. In the present study, MS Binding Assays utilizing UNC0642 as a reporter ligand have been established. Thus, the binding of UNC0642 towards *Torpedo*-nAChR has been characterized in MS saturation and competition experiments. According to the results, UNC0642 addresses the MB327 binding site of the *Torpedo*-nAChR. This conclusion is further supported by the outcome of *ex vivo* studies performed with poisoned rat diaphragm muscles as well as by *in silico* studies predicting the binding mode of the most affine analog UNC0646 in the recently proposed binding site of MB327 (MB327-PAM-1). The new MS Binding Assays based on the commercially available reporter ligand UNC0642 as one of the most affine ligands for the MB327 binding site are a potent and valuable alternative to established assays.

## 1. Introduction

The poisoning with organophosphorus compounds (OPCs) as a result of exposure to respective insecticides or nerve agents represents a severe health problem (Buckley et al., 2004; Costanzi et al., 2018; Eddleston and Phillips, 2004; John et al., 2018). If not treated properly, an intoxication with OPCs can culminate in a cholinergic crisis, which can finally lead to death because of respiratory failure (Holmstedt, 1959; Newmark, 2004). As the principal mode of action, all OPCs have in common to inactivate acetylcholinesterase (AChE), an enzyme that is in charge of the breakdown of the neurotransmitter acetylcholine in the synaptic cleft of cholinergic neurons. This results in the accumulation of this neurotransmitter in the synaptic cleft whereupon the corresponding receptors, the muscarinic acetylcholine receptor (mAChR) and the nicotinic acetylcholine receptor (nAChR), become overstimulated. Being the main cause for the OPC-induced adverse health effects, medical measures aim, in general, at the elimination of this overstimulation. Thus, the standard treatment of OPC intoxication includes the application of the mAChR antagonist atropine to reduce neuronal signaling mediated by this receptor and the use of oximes, such as obidoxime, to reactivate the AChE and, thus, to lower the acetylcholine level by enzymatic breakdown (Shih et al., 2007; Thiermann and Worek, 2022; Worek et al., 2005). The use of oximes has, however, often been found to be insufficiently effective, which strongly depends on the type of OPC causing the intoxication (Thiermann et al., 2016). Hence, there is a strong need for pharmacological agents that may counteract overstimulation and resulting desensitization of the receptors by direct intervention at nAChRs.

Though antagonists for the acetylcholine binding site of the nAChR are known, their application to reduce nAChR overstimulation – in analogy to that of atropine for mAChR – is not feasible, as the therapeutic window of these compounds is too small (Sheridan, 2005). Yet, as an alternative, pharmacological agents that interact with nAChRs via an allosteric binding site and restore the functional activity of desensitized receptors may be applied. Among non-oxime bispyridinium salts, a series of compounds has been identified that act this way, of which MB327 (Figure 1) can be considered the most prototypic representative. Interestingly, in electrophysiological measurements, MB327 has been demonstrated to restore the functional activity of nAChRs, which had been desensitized by overstimulation with orthosteric ligands (Niessen et al., 2016; Niessen et al., 2018). Furthermore, *in silico* studies led to the identification of a potential allosteric binding pocket for MB327 at nAChRs, termed MB327-PAM-1 binding site, where MB327 is thought to act as an allosteric modulator reestablishing receptor function (Kaiser et al., 2023). In addition, pharmacological effects for MB327 and some analogs have been recorded that demonstrate that these compounds can restore the muscle force of rat diaphragm muscles defunctionalized by soman treatment in *ex vivo* experiments (Niessen et al., 2018; Seeger et al., 2012). The ability of MB327 to reactive soman-poisoned intercostal muscles has also been shown for tissue from humans (Seeger et al., 2012). Moreover, MB327 (respectively the corresponding methanesulfonate salt) was found to increase the survival rate of nerve agent-poisoned guinea pigs in *in vivo* studies when applied as a drug agent (Timperley et al., 2012; Turner et al., 2011). Unfortunately, with its low potency and small therapeutic window, MB327 is far from fulfilling the requirements for a drug candidate (Kassa et al., 2022). Hence, great efforts have been undertaken to identify more potent allosteric modulators of the MB327 binding site.

**Figure 1.**
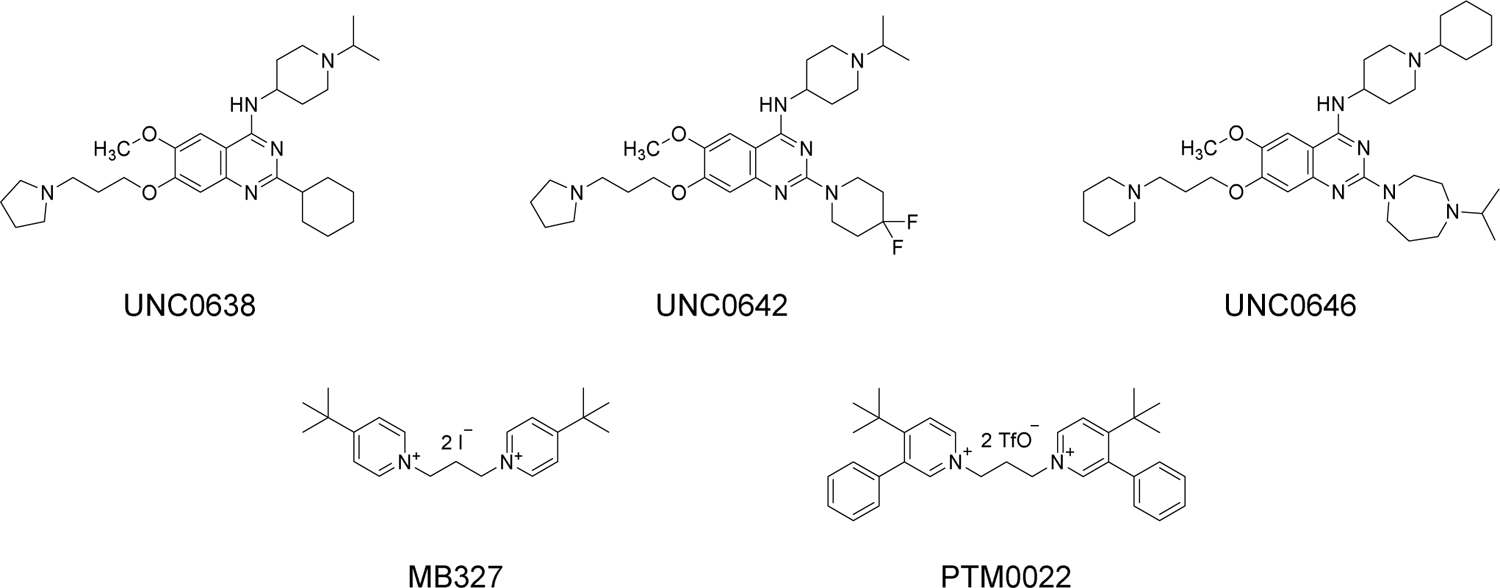
Ligands of the MB327 binding site of the nAChR: MB327, PTM0022, UNC0638, UNC0642, and UNC0646.

Sichler et al. have developed an MS Binding Assay addressing the MB327 binding site using [^2^H_6_]MB327 as a reporter ligand (Sichler et al., 2018). This has been extensively used for the characterization of the binding affinities of a plethora of bispyridinium salts related to MB327 that had been synthesized to gain insight into the structure-activity relationship of this compound class and to finally unveil representatives with distinctly higher affinities than MB327. Although binding affinities of most compounds were in the range of that of MB327 (p*K*_i_ = 4.73 ± 0.03), one bispyridinium salt, PTM0022, delineated from MB327 by two additional phenyl residues, was found to surpass the binding affinity of MB327 to a small but statistically significant extent (p*K*_i_ = 5.16 ± 0.07) (Rappenglück et al., 2018a, b). Significant progress was finally achieved when Sichler et al. used their [^2^H_6_]MB327-based MS Binding Assays addressing the MB327 binding site of the nAChR for the screening of a commercial compound library (Sichler et al., *submitted*). That way, a group of quinazoline derivatives with high affinities for the MB327 binding site was identified, with the highest affinities displayed by UNC0638 (p*K*_i_ = 6.01 ± 0.10), UNC0642 (p*K*_i_ = 5.97 ± 0.05), and UNC0646 (p*K*_i_ = 6.23 ± 0.02) (Figure 1).

The present study aimed to develop new MS Binding Assays for the MB327 binding site of the nAChR utilizing one of these quinazoline derivatives as reporter ligand. Such binding assays were expected to be more robust and specific due to the distinctly increased affinity of the employed reporter ligand as compared to that of MB327 in the [^2^H_6_]MB327-based MS Binding Assays. Moreover, such binding assays should provide additional solid information on the interaction of the aforementioned quinazoline derivatives with the MB327 binding site of the nAChR, which so far had only been studied in competitive [^2^H_6_]MB327 MS binding experiments. For the new binding assays, the concept of MS Binding Assays was followed due to simple reasons: As often discussed in the literature, this type of binding assays benefits from not requiring radiolabeled substances, which makes them highly flexible with regard to the compounds used as reporter ligands. Besides, MS Binding Assays do not suffer from drawbacks that commonly arise when radioactivity is involved in an experimental setting (Höfner and Wanner, 2015; Wanner et al., 2007).

To gain knowledge on the intrinsic activity of the newly identified quinazoline derivatives addressing the MB327 binding site, their capability to restore muscle force in *ex vivo* experiments with soman-poisoned diaphragm muscle tissues were studied, too. In addition, using docking approaches and molecular dynamics (MD) simulations, the binding mode and key interaction partners of the most affine analog UNC0646 in MB327-PAM-1 were studied.

## 2. Results and Discussion

### 2.1 MS Binding Assays addressing the *Torpedo*-nAChR with quinazoline derivatives

#### 2.1.1 LC-ESI-MS/MS method development

For performing the MS Binding Assays with one of the newly identified ligands of the MB327 binding site with a quinazoline scaffold, i.e., UNC0638, UNC0642, and UNC0646, first, a reliable LC-MS/MS method for quantification was needed. To this end, a triple quadrupole mass spectrometer in the multiple reaction monitoring (MRM) mode, in combination with a pneumatically assisted electrospray ionization source (ESI) and an HPLC system, should be used. This setup has repeatedly been demonstrated to achieve the selectivity and sensitivity required for marker quantification in MS Binding Assays and, thus, to be well suited for this purpose (Ackermann et al., 2021; Ackermann et al., 2019; Grimm et al., 2015; Hess et al., 2011; Neiens et al., 2018; Neiens et al., 2015). In the literature, for the three compounds UNC0638, UNC0642, and UNC0646, only MS studies reporting their parent ions but no mass fragmentations are known (Liu et al., 2011; Liu et al., 2013; Vedadi et al., 2011). Accordingly, first, the mass transitions of UNC0638, UNC0642, and UNC0646 were analyzed in direct infusion experiments. The most intense mass transitions found, all of which originated from the parent ion [M+H]^+^, were (see Figure S1 for product ion spectra): UNC0638 *m*/*z* 510.3/112.2, UNC0642 *m*/*z* 547.3/112.1, and UNC0646 *m*/*z* 622.5/126.1.

Next, a suitable LC-ESI-MS/MS method for the quantification of these compounds had to be developed. To enable a reasonably high sample throughput of the MS Binding Assays, such a method should have a short run time while still separating the analyte from contents in the sample matrix interfering with the MS analysis, which, according to our experience, can commonly be reached when the retention factors of the analytes are > 1. In their recent work, Sichler et al. described LC-ESI-MS/MS quantification methods for MS Binding Assays that were all based on the same LC conditions, though the analytes, all polar ligands, varied (e.g., MB327 and phencyclidine) (Sichler et al., 2018). This method is based on a YMC-Triart Diol-HILIC column operated under classical HILIC conditions [mobile phase acetonitrile/ammonium formate buffer (20 mM, pH 3.0) = 80:20; flow rate 800 µL/min]. The quinazoline derivatives UNC0638, UNC0642, and UNC0646 are to be expected to be protonated and, consequently, to possess a high polarity under these chromatographic conditions. Hence, we reasoned that these conditions might also be suitable for their analysis. Indeed, the chromatograms for UNC0638 and UNC0642 obtained with these LC-parameters were satisfying, with the retention factor *k* amounting to 1.7 and 2.1 for UNC0642 and UNC0638, respectively, and the run time to < 3 min (for a chromatogram, see Figure 2). Under the same chromatographic conditions, UNC0646, however, yielded a peak with a retention factor of > 10, which was far too high for the intended purpose and the peak shape was poor. This issue could be overcome by raising the amount of buffer in the mobile phase from 20% to 35%, whereas the flow rate had to be reduced from 800 to 600 µL/min to not exceed the limits for back pressure. This led to a retention factor *k* = 1.2 for UNC0646 (for a chromatogram, see Figure S2), which was in the desired range. Finally, preliminary binding experiments should be performed to explore, which of the three compounds might be best suited as a reporter ligand for the planned MS Binding Assays.

**Figure 2.**
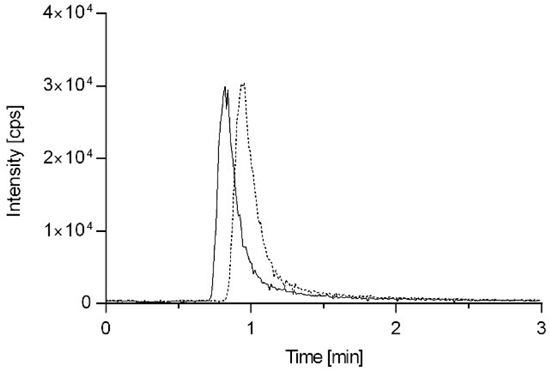
LC-ESI-MS/MS-MRM chromatogram of a matrix standard containing the reporter ligand UNC0642 (solid line) at a concentration of 5 nM and the internal standard UNC0638 (dashed line) at 10 nM. For quantification, the mass transitions *m*/*z* 547.3/112.1 and *m*/*z* 510.3/112.2 for UNC0642 and UNC0638, respectively, were used. A YMC-Triart Diol-HILIC (50 mm x 2.0 mm, 3 µm) column was used as a stationary phase in combination with an 80:20 (*v*/*v*) mixture of acetonitrile and ammonium formate buffer (20 mM, pH 3.0) as mobile phase. The injection volume amounted to 10 µL and the flow rate to 800 µL/min.

#### 2.1.2 Preliminary binding experiments and determination of final assay conditions

When developing a binding assay, irrespective of whether this is, e.g., a radioligand or MS Binding Assay, a decision regarding the technique has to be made that is used for the separation of the target protein with the bound reporter ligand from the rest of the incubation mixture containing the non-bound ligand. In general, filtration is preferred for separation, as it offers an efficient way of handling binding samples and a high sample throughput. Unfortunately, for a filtration-based binding assay, a ligand with a high binding affinity is required. According to the literature, the *K*_d_ value should be in the range of 10^-7^ to 10^-8^ M or lower (Hulme and Trevethick, 2010; McKinney and Raddatz, 2006), as otherwise the *k*_off_ rate (indirectly reflected by the *K*_d_ value) is too high and the loss of specifically bound reporter ligand during washing steps will exceed the 10% limit, which will affect the results to a non-tolerable extent. As an alternative to the separation process, centrifugation may be used for ligands with affinities that are too low for filtration. This approach suffers, however, from the fact that the separation step is laborious and the throughput rather low. From the p*K*_i_ values determined in the [^2^H_6_]MB327 MS Binding Assay for UNC0638 (p*K*_i_ = 6.01 ± 0.10), UNC0642 (p*K*_i_ = 5.97 ± 0.05), and UNC0646 (p*K*_i_ = 6.23 ± 0.02), it is obvious that with these compounds as reporter ligands only centrifugation can be used for the separation step. As UNC0646 has the highest affinity for the MB327 binding site compared to UNC0638 and UNC0642, it was first chosen as a reporter ligand. When following the general assay procedure Sichler et al. had developed for the centrifugation-based MS Binding Assay with [^2^H_6_]MB327 as reporter ligand addressing *Torpedo*-nAChR, we were indeed able to observe specific binding for UNC0646 in preliminary binding experiments (Rappenglück et al., 2018b; Sichler et al., 2018). For the sake of completeness, also filtration was tested as a separation technique with UNC0646 as the reporter ligand. The experiments, however, led to results suggesting that most of the target-bound ligand had been lost during this assay based on the filtration approach. Hence, centrifugation should be used for the separation step for all further experiments.

Much to our surprise, when attempting to finalize the conditions for the MS Binding Assay with UNC0646 as the reporter ligand, we encountered repeatedly difficulties regarding the reproducibility of the compound quantification. Hence, we decided to test UNC0642 as a reporter ligand in MS Binding Assays, although its binding affinity is lower than that of UNC0646.

Again, for the MS Binding Assays with UNC0642 as a reporter ligand, we kept close to the conditions that Sichler et al. had established for their centrifugation-based [^2^H_6_]MB327 MS Binding Assay addressing the *Torpedo*-nAChR (see Experimental Section for details). However, in the case of the new binding assay, non-specific binding was determined by the competitor approach, whereas Sichler et al. had applied the heat shock method, by which the target material is denatured to lose specific binding (Sichler et al., 2018). For the new binding assays, highly affine ligands of the MB327 binding site were available that appeared well suited for the determination of non-specific binding by the competitor approach with competitors not yet identified when Sichler et al. developed their [^2^H_6_]MB327 MS Binding Assay. Hence, we opted for this approach, as this is the most common one (Hulme and Trevethick, 2010; Motulsky and Neubig, 2002). As a competitor for the determination of non-specific binding, UNC0646 was selected as one of the two quinazoline derivatives with high affinities for the MB327 binding site, UNC0638 and UNC0646. This decision was made, as the reporter ligand should be quantified using an internal standard to improve the robustness of the quantification method. Hence, one of the two aforementioned quinazoline derivatives was needed to this end. As only UNC0638 exhibits a chromatographic behavior similar to that of the reporter ligand UNC0642 but not UNC0646 (see previous chapter), which is essential for a compound to be used as an internal standard, UNC0638 could serve this function. Accordingly, for determining non-specific binding in the binding assay, quinazoline derivative UNC0646 had to be used.

#### 2.1.3 LC-ESI-MS/MS method and method validation

With the conditions of the MS Binding Assays and, thus, also the matrix of the analytical samples being defined, the preconditions for the validation of the LC-ESI-MS/MS method for the quantification of the reporter ligand UNC0642 with UNC0638 as internal standard were given. As indicated above, the LC method was largely the same as the one developed by Sichler et al. (Sichler et al., 2018). In Figure 2, a chromatogram of the ligand UNC0642 and the internal standard UNC0638 obtained applying these LC conditions is given with the most important parameters of the LC-ESI-MS/MS method being listed in the caption (for further details see Experimental Section).

For the validation of the analytical method, the recommendations of the FDA guidance for bioanalytical method validation were followed regarding the criteria linearity, accuracy, precision, sensitivity, and selectivity (FDA, 2018). The results of the validation process are briefly discussed in the following. Detailed validation data of the corresponding three validation series can be found in the SI (see Figure S3 and Table S1).

For the validation experiments, matrix blank and matrix zero calibrator samples were prepared in analogy to the samples of the MS binding experiments. Matrix zero calibrator samples were used to create calibration standards and quality control samples (see Experimental Section for details). Calibration standards were prepared for eight different concentrations in the range from 50 pM (lower limit of quantification, LLOQ) to 75 nM and quality control samples for the four concentrations 50 pM (LLOQ), 500 pM, 5 nM, and 50 nM. In calibration standards and quality control samples, the internal standard UNC0638 was present at 10 nM.

To evaluate the linearity of the quantification method in the investigated concentration range (50 pM to 75 nM), the calibration standards were analyzed via linear regression to obtain a calibration curve (see Experimental Section for details). The criteria of the FDA guideline for linearity demand calibration standard deviations from the nominal concentrations to be within a limit of ± 15% (± 20% at LLOQ). As we determined calibration standard deviations from the nominal concentrations in the range from 93% to 112%, the criteria of the FDA guideline for linearity were fulfilled. Deviations of the measured concentrations from nominal concentrations within ±15% (± 20% at LLOQ) are required for accuracy by the FDA guidelines for quality control samples. For intra-run samples, accuracies from 91 - 103% and for inter-run samples accuracies from 94 - 101% were found, thus fulfilling the acceptance criteria of the FDA guideline for these criteria. The quality control samples were also examined regarding precision (expressed by the relative standard deviation), which amounted to 2.5 - 6.5% and 4.0 - 6.3% for intra-run and inter-run precision, respectively, being in line with the acceptance criteria of ± 15% (± 20% at LLOQ) of the FDA guidelines. Also, the sensitivity of the LC-ESI-MS/MS method was guaranteed as the intensity of the peak corresponding to the LLOQ of 50 pM as compared to the noise signals was in line with the required signal-to-noise ratio of at least five. When a matrix blank sample was measured, no interference was found, demonstrating the selectivity of the established LC-ESI-MS/MS method. Overall, all studied validation criteria of the analytical method comply with the standards defined by the FDA guidelines.

#### 2.1.4 UNC0642 MS Binding Assays

### Saturation Experiments

Next, with the validated quantification method for UNC0642 at hand, the binding of this compound to *Torpedo*-nAChR should be characterized in saturation experiments. For saturation experiments, the target is in general incubated with the reporter ligand in a concentration range from 0.1 *K*_d_ to 10 *K*_d_ for the determination of total binding. As the *K*_d_ of UNC0642 was expected to be in the very low micromolar range, we investigated fifteen different reporter ligand concentrations ranging from 200 nM to 100 µM for total binding. For the determination of non-specific binding, a further set of binding samples containing UNC0646 as competitor was prepared. Because of its low solubility, UNC0646 could only be used in an assay concentration of up to 100 µM. Since the affinity of UNC0646 (p*K*_i_ = 6.23 ± 0.02) determined in [^2^H_6_]MB327 MS Binding Assays is similar to that of the reporter ligand UNC0642 (p*K*_i_ = 5.97 ± 0.05), according to common rules, at least a hundredfold excess of the competitor UNC0646 over the reporter ligand UNC0642 had to be applied. Hence, because of the limited solubility of UNC0646, non-specific binding could only be measured for the five lowest reporter ligand concentrations (200 nM - 1 µM). Based on these data, a linear regression function, which was forced through zero, was established and finally used for the calculation of non-specific binding values for all reporter ligand concentrations employed in the assay (Davenport and Russell, 1996).

Specific binding as the difference between total and non-specific binding was finally analyzed by non-linear regression, generating saturation isotherms that revealed a binding affinity for UNC0642 of 6.7 ± 0.4 µM (*K*_d_) and a maximum density of binding sites *B*_max_ = 2980 ± 130 pmol/mg protein. The results of a representative saturation experiment are depicted in Figure 3. The obtained *K*_d_ value of 6.7 ± 0.4 µM, corresponding to a p*K*_d_ of 5.17 ± 0.03, is in reasonable accordance with the p*K*_i_ value of 5.97 ± 0.05 previously determined by Sichler et al. (Sichler et al., *submitted*). The *B*_max_ value found in our experiments appears to be rather high, which may be explained by recent *in silico* experiments suggesting that there are multiple MB327 binding sites in the nAChR (Kaiser et al., 2023). Overall, the results of these saturation experiments indicate that UNC0642 binds to nAChR in a specific and saturable manner, which in combination with the data found by Sichler et al. in competition experiments with [^2^H_6_]MB327 as reporter ligand (Sichler et al., *submitted*), further supports the assumption that both address the same binding site, the MB327 binding pocket of *Torpedo*-nAChR.

**Figure 3.**
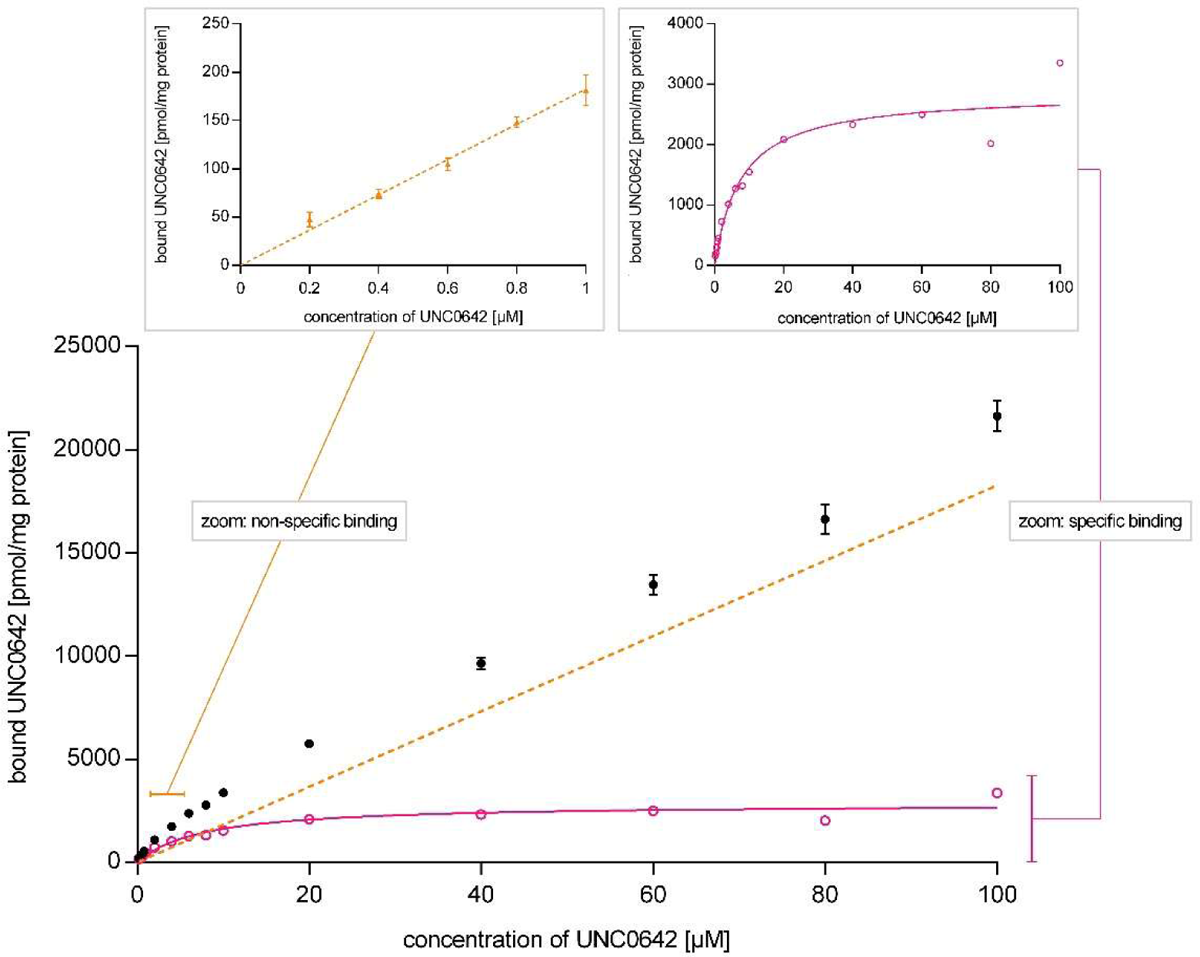
Representative saturation experiment for UNC0642 binding to *Torpedo*-nAChR. Total binding (black circles) and non-specific binding (orange triangles). Linear regression of non-specific binding is shown as an orange dashed line. Specific binding (pink circles) was calculated as the difference between total binding and non-specific binding and analyzed by non-linear regression (solid pink line). Experimental values are means ± SD, n = 3.

### Competition experiments

Finally, with the methodology for performing saturation experiments with UNC0642 as a reporter ligand at hand, competitive MS Binding Assays addressing the MB327 binding site of *Torpedo*-nAChR should be established. These should be used to characterize the binding affinities of a representative set of ligands of the MB327 binding pocket known from [^2^H_6_]MB327 MS Binding Assays. As the binding affinity of MB327 towards the MB327 binding site of *Torpedo*-nAChR is rather low, the results from MS Binding Assays based on [^2^H_6_]MB327 as a reporter ligand might deviate from the real value. Yet, these results should still be a reasonable basis for a comparison with the data obtained from the new MS Binding Assay with UNC0642 as a reporter ligand. The comparison might allow us to validate the results of the UNC0642 MS Binding Assay and, in addition, further support the assumption that UNC0642 and MB327 address the same binding pocket of *Torpedo*-nAChR.

The set of competitors to be studied in the UNC0642 MS Binding Assays contains MB327, as the reporter ligand from the [^2^H_6_]MB327 MS Binding Assay and most prototypic representative of bispyridinium salts addressing nAChR, and PTM0022, as this compound shows the highest affinity so far found within the class of bispyridinium salts (Rappenglück et al., 2018b). Furthermore, the quinazoline derivative UNC0646 should be included in this set of test compounds, as it represents the compound with the highest affinity for the MB327 binding site so far known (Sichler et al., *submitted*). For control purposes, finally, also the effect of carbachol, which is a well-known ligand of the orthosteric binding site of nAChR, in the new competitive MS Binding Assays with UNC0642 as the reporter ligand should be studied.

The competition experiments were performed in analogy to the saturation experiments with the following adaption. The concentration of the reporter ligand UNC0642 was kept constant at 1 µM. Binding samples were provided with increasing concentrations of the respective test compounds, usually covering a range of three orders of magnitude around the expected IC_50_. After quantification of the reporter ligand via LC-ESI-MS/MS, the obtained data was normalized. For this, binding samples had been prepared, which contained no competitor (equivalent to 100% specific binding) or 100 µM UNC0646 as competitor (i.e., non-specific binding, 0% specific binding). Competition curves were created by non-linear regression yielding the respective IC_50_ values of the test compounds from which *K*_i_ values were calculated according to the Cheng-Prusoff equation (see Experimental Section for details).

The competition curves that resulted when the above-mentioned test compounds were characterized in the new competitive MS Binding Assays with UNC0642 as reporter ligand – in three independent experiments in every case – are given in Figure 4a and 4b.

**Figure 4.**
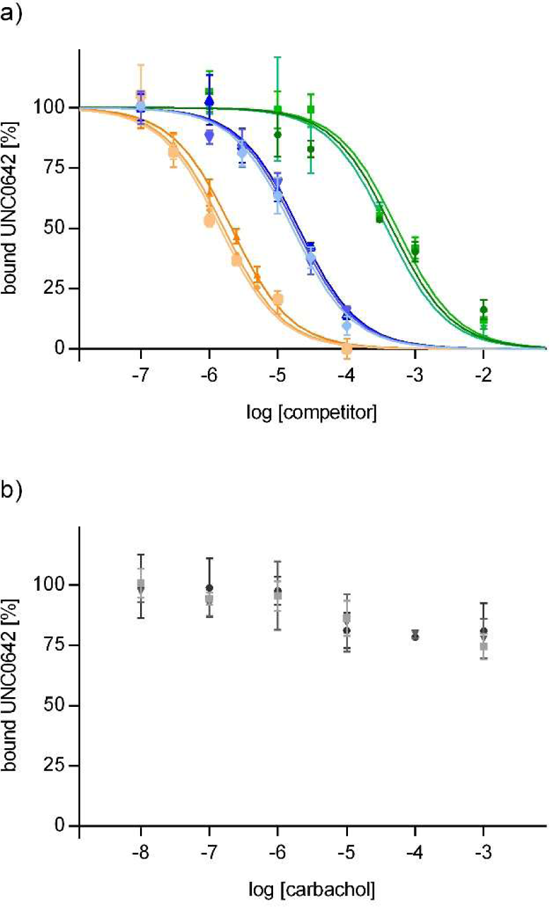
a) Competition curves obtained for MB327 (green), PTM0022 (blue), and UNC0646 (orange). b) Influence of carbachol on UNC0642 binding. Data points (mean ± SD, *n* = 3) represent the specific binding of UNC0642.

Except for carbachol, for all other test compounds, the shape of the competition curves was well in line with theoretical models. The analysis of the curves revealed IC_50_ values from which p*K*_i_ values of 3.40 ± 0.04, 4.80 ± 0.03, and 5.83 ± 0.05 for MB327, PTM0022, and UNC0646, respectively, were calculated. Overall, p*K*_i_ values found in the new MS Binding Assay utilizing UNC0642 as reporter ligand were lower than those determined in the [^2^H_6_]MB327 MS Binding Assay, but most importantly the rank order of affinities remained the same (MB327 < PTM0022 < UNC0646, see Table 1). Interestingly, also carbachol affected the binding of UNC0642. Up to a concentration of 1 µM carbachol, the UNC0642 binding remained largely unchanged, whereas it decreased for higher carbachol concentrations to reach a plateau of 75 - 80% at about 10 µM, which persists up to the highest concentration applied (1 mM). Notably, the change of UNC0642 binding occurs in the same range of carbachol concentration – 1 µM to 10 µM – that is known from functional studies to affect the transition of *Torpedo*-nAChR from its resting into its active state (Niessen et al., 2016). If the concentration of an orthosteric ligand present at the nAChR is far above the amount required for activation, the receptor switches into a desensitized state (Papke, 2014). This has to be taken into account for the analysis of the above-described data, as the transition of *Torpedo*-nAChR into the desensitized state likely occurs at a carbachol concentration covered in the experiments, i.e., ≥ 100 µM (Währa et al., 2023). Thus, the decrease of UNC0642 binding to the MB327 binding site upon increasing the carbachol concentration is likely the result of a conformational change due to the orthosteric ligand binding, which exerts an allosteric effect between the orthosteric and the MB327 binding site (Kaiser et al., 2023). However, for a better understanding of the interaction between the MB327 binding site and the orthosteric binding site, further studies are needed.

**Table 1.** p*K*_i_ values obtained from IC_50_ values determined in UNC0642 MS Binding Assays and [^2^H_6_]MB327 MS Binding Assays, respectively (Rappenglück et al., 2018b; Sichler et al., *submitted*).

| Compound | UNC0642<br>MS Binding Assay | [$^2\text{H}_6$]MB327<br>MS Binding Assay |
| --- | --- | --- |
| | $pK_i$ | |
| MB327 | $3.40 \pm 0.04$ | $4.73 \pm 0.03$ |
| PTM0022 | $4.80 \pm 0.03$ | $5.16 \pm 0.07$ |
| UNC0646 | $5.83 \pm 0.05$ | $6.23 \pm 0.02$ |

Overall, according to the results of the UNC0642 MS Binding Assays, it is reasonable to conclude that UNC0642 alike MB327 addresses the MB327 binding site of the nAChR. In particular, the binding of UNC0642 can completely be inhibited by MB327, and both the MS Binding Assay based on UNC0642 and on [^2^H_6_]MB327 yield p*K*_i_ values that are in reasonable to good agreement and lead to the same rank order of potencies in competitive experiments for the set of ligands studied (see Table 1). Hence, the UNC0642 MS Binding Assays represent a valuable new tool for the characterization of the affinity of ligands of the MB327 binding site of the nAChR. With the binding affinity of UNC0642 being distinctly higher than that of [^2^H_6_]MB327, the reporter ligands of the UNC0642 and [^2^H_6_]MB327 MS Binding Assays, the former can be considered more robust with regard to its performance and results than the latter. In addition, the former MS Binding Assay has the advantage that its reporter ligand, UNC0642, is commercially available, which eases its setup.

### 2.2 *In silico* investigation of the UNC0646 binding mode in MB327-PAM-1

Recently, we proposed a novel binding site, MB327-PAM-1, in nAChR for binding of MB327 that can explain the allosteric modulation relevant for treating poisoning with OPC (Kaiser et al., 2023). MB327-PAM-1 is located in between two adjacent subunits at the transition of the extracellular to the transmembrane region and is different from two allosteric and one orthosteric binding pocket that had been proposed before for bispyridinium compounds using *in silico* methods (Epstein et al., 2021; Wein et al., 2018). To investigate the binding site of the most-affine ligand in this study, UNC0646, first, we performed flexible docking experiments to place the ligand in MB327-PAM-1. To do so, we selected all residues within 9 Å of the centrally located E199α (respectively, Q209β, E210δ, E200γ in the other subunits) as a potential binding site. UNC0646 was placed similarly at the negative side of the two α subunits, whereas in the other three subunits, the ligand was either placed in the middle between two possible binding sites or more towards the pore, which would result in a high solvent exposure (SI Figure S4). In between the γ- and α-subunit (binding site A), the best-scored conformation contains a twist conformation of the cyclohexane ring and a boat conformation of the piperidine ring of the side chain in the 4-position of the quinazoline ring (SI Figure S5). As these ring conformations are energetically unfavorable, we chose the second best-scored conformation. The orientation of UNC0646 in the binding pocket is similar there but the rings have chair conformations. Overall, in both binding sites at the negative side of the α-subunits, the orientation of UNC0646 is comparable. However, while in between the β- and α-subunit (binding site B) the nitrogen of the piperidine ring in the 4-position of the quinazoline ring is interacting with E199α, the ligand is placed more deeply in the binding site A, which facilitates interaction between this nitrogen and E65γ (Figure 5A, SI Figure S6).

**Figure 5.**
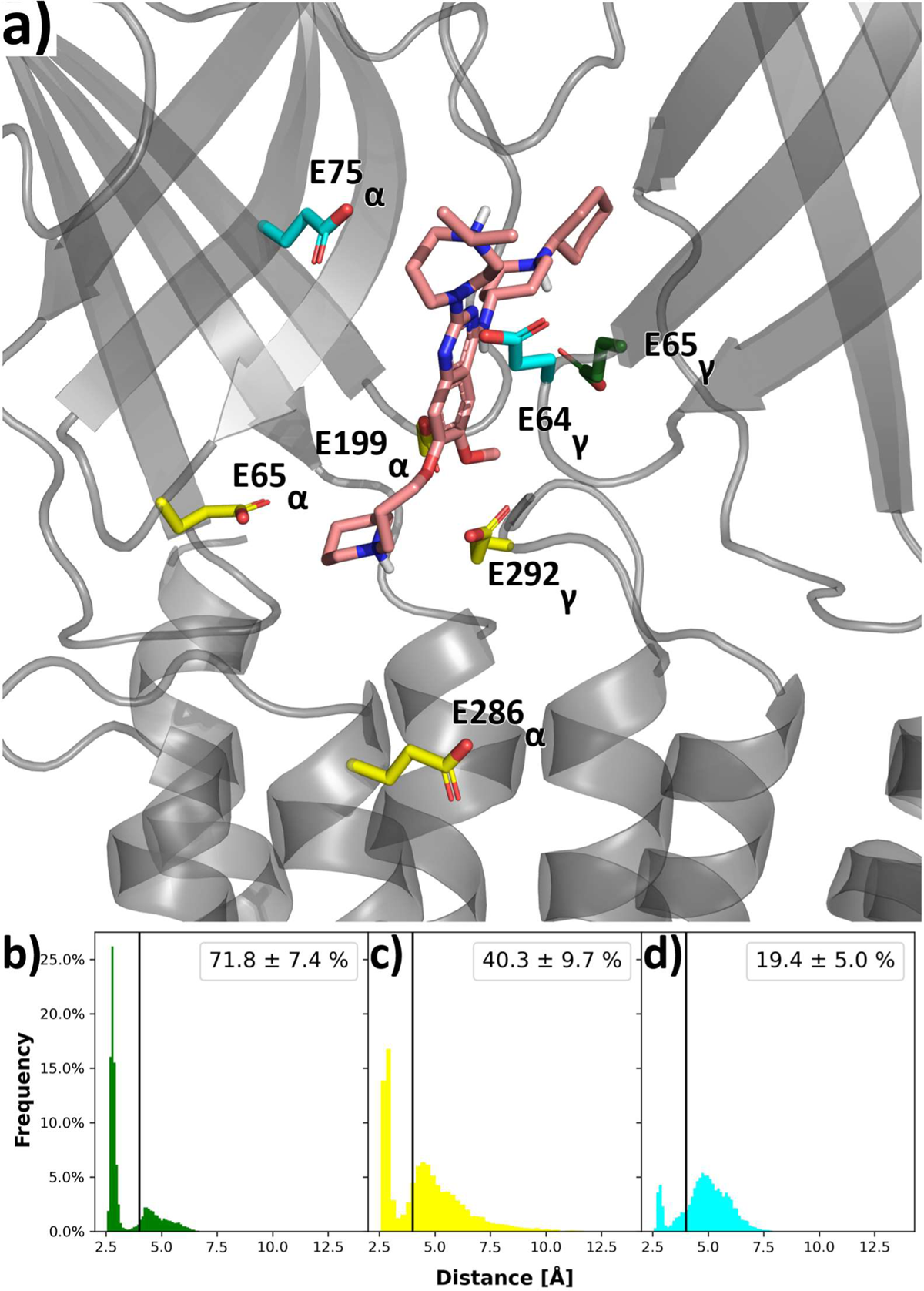
UNC0646 and interacting residues in *Torpedo* nAChR. a) Docked binding mode of UNC0646 as starting point for MD simulations. b) Minimal distance of the positively charged piperidyl nitrogen in the sidechain at position 4 to the carboxylate oxygens of E65γ. c) Minimal distance of the positively charged piperidyl nitrogen in the side chain at position 7 to the carboxylate oxygens of E65α, E199α, E286α, and E292γ. d) Minimal distance of the positively charged nitrogen in the diazepane ring to the carboxylate oxygens of E75α and E64γ. Amino acids in panel A are colored according to the plot colors in panels b) - d). The values in panels B-D indicate the mean ± SEM (taken over 12 replicas each) of the frequency of hydrogen bonds (distance of nitrogen to carboxylate oxygen < 4 Å).

To further scrutinize interactions with surrounding amino acids, we performed 12 replicas of 500 ns long unbiased MD simulations starting from the docked conformations of UNC0646 in binding sites A and B resulting in 6 μs (12 x 500 ns) of cumulative simulation time. During MD simulations, the receptor and membrane remained structurally virtually invariant (SI Figure S7, S8). Throughout the MD simulations, UNC0646 showed smaller movements in binding site A (RMSD = 3.02 ± 0.30 Å) than binding site B (RMSD = 5.13 ± 0.58 Å, *p* = 0.004 according to a two-sided t-test). Furthermore, UNC0646 leaves the binding site B in six out of 12 replicas (distance to I65α < 5 Å in the last frame, as done previously (Kaiser et al., 2023)), whereas this does not occur in any of the 12 replicas in binding site A. Together, this suggests that the orientation of UNC0646 in binding site A is preferred.

Thus, we used this orientation to further predict important residues for interactions, in particular, salt bridge interactions of the three tertiary amines in the substituents of the quinazoline ring with the glutamates in the binding site; glutamates were ranked as the most important residues for ligand binding in per-residue decompositions of the effective binding energy computed with MMPBSA (SI Figure S9). E65γ, previously described to be important for interactions with MB327 (Kaiser et al., 2023), shows the most conserved interactions with the piperidyl moiety in position 4 of the quinazoline ring (71.8 ± 7.4% of all frames, Figure 5). Second, the positively charged nitrogen of the substituent in the 7-position interacts primarily with E69α (28.2 ± 10.5%). As this nitrogen is located in an area surrounded by four glutamates (E69α, E199α, E286α, and E292γ), additional salt bridge interactions can form. Considering all four glutamates, the nitrogen is interacting with carboxylate oxygens in 40.3 ± 9.7% of all frames. In contrast, the tertiary amine nitrogen in the diazepane ring at position 2 shows only minor interactions with the two surrounding glutamates (E75α, and E64γ; in 19.4 ± 5.0% of all frames). These results indicate that the side chain in the 4-position of the quinazoline ring is most important for forming salt bridge interactions whereas the diazepane ring is least important. These findings are in line with a structure-affinity relationship deduced from compounds UNC0646, UNC0642, and UNC0638, where removing the positive charge in the 2-position of the quinazoline ring has only minor effects on ligand affinity. Furthermore, the findings are supported by residue conservation analysis according to which E65γ, E69α, E199α, and E286α are highly conserved among different subunits of the *Torpedo* and human muscle-type nAChR such that acidic side chains at each position are available in at least three of the five subunits and hydrogen bond acceptors are available in all subunits but one (Table 2). By contrast, E292γ, E75α, and E64γ are less conserved (Table 2).

**Table 2.** Sequence similarity of *Torpedo* and human adult muscle-type nAChR subunits. Amino acids shown in Figure 5 are represented with green shadings. Amino acids at structurally homologous positions in other subunits are shown on a white background; amino acids with deviating physicochemical properties are shown in italics.

| Human muscle-type nAChR |  |  |  | Torpedo nAChR |  |  |  |
| --- | --- | --- | --- | --- | --- | --- | --- |
| $\alpha$ | $\beta$ | $\delta$ | $\epsilon$ | $\alpha$ | $\beta$ | $\delta$ | $\gamma$ |
| E | E | E | E | E69 | E | E | E |
| *T* | *S* | *T* | *T* | E75 | *T* | *T* | *T* |
| E | *Q* | E | E | E199 | *Q* | E | E |
| E | D | *K* | *Q* | E286 | D | *Q* | *Q* |
| *N* | D | E | E | *N* | *I* | D | E64 |
| *Q* | E | E | E | *Q* | E | E | E65 |
| *S* | E | *A* | E | *S* | E | E | E292 |

Based on a representative binding mode of UNC0646 during MD simulations in binding site A, we replaced the substituents of the quinazoline ring of UNC0646 to match those of UNC0642 using MOE and subsequently minimized the ligand in the binding site (SI Figure S10). The substituents in 4- and 7-position of the quinazoline ring of UNC0642 interact similarly as those of UNC0646, whereas due to a lack of protonation sites interactions with the ring system in 2-position are missing.

### 2.3 Evaluation of the muscle force recovery of quinazoline-based compounds

To gain more knowledge on the intrinsic effects of UNC0638, UNC0642 and UNC0646 we performed myographic experiments with soman-poisoned and also un-poisoned rat diaphragms. The corresponding results of these experiments are summarized in Figure 6. Interestingly, the compound with the highest known affinity to the MB327 binding site, UNC0646, was the only compound in this series of experiments that did not seem to have a beneficial effect on the restoration of muscle force after soman poisoning. UNC0638 and UNC0642 instead induced the regeneration of muscle force at a maximum of 30 µM and 10 µM, respectively. The highest extent of recovery was observed for a stimulation frequency of 20 Hz and amounted to 18.4 ± 16.1% for UNC0638 at 30 µM and 16.2 ± 12.8% for UNC0642 at 10 µM (mean ± SD, n = 5). The maximum amplitudes of UNC0638 and UNC0642 are thus lower than the maximum amplitude observed for MB327 [approximately 30% at 20 Hz (Niessen et al., 2018; Seeger et al., 2012)], but favorably the concentration needed to generate the described effect was distinctly lower for UNC0638 and UNC0642 than for MB327, which showed its maximum effect at a concentration of 300 µM. To obtain a recovery comparable to that exhibited by UNC0638 and UNC0642 at a concentration of 30 µM and 10 µM, respectively, MB327 had to be used at 100 µM (Niessen et al., 2018; Seeger et al., 2012). Noteworthy, the muscle force restoration effected by UNC0638 and UNC0642 declined at higher concentrations after the maximum had been reached at 30 µM and 10 µM, respectively. This phenomenon has been observed for MB327 and some MB327 analogs before (Niessen et al., 2018). It has been speculated, that counteracting effects, mediated by different binding sites may be responsible for the observed course of muscle force as a function of the compound concentration. In the present case, i.e. for UNC0638, UNC0642, and UNC0646, this theory is supported by results, that have been obtained in myographic experiments performed in analogy to those above except for using native, functionally active instead of soman-poisoned rat-diaphragms.

**Figure 6.**
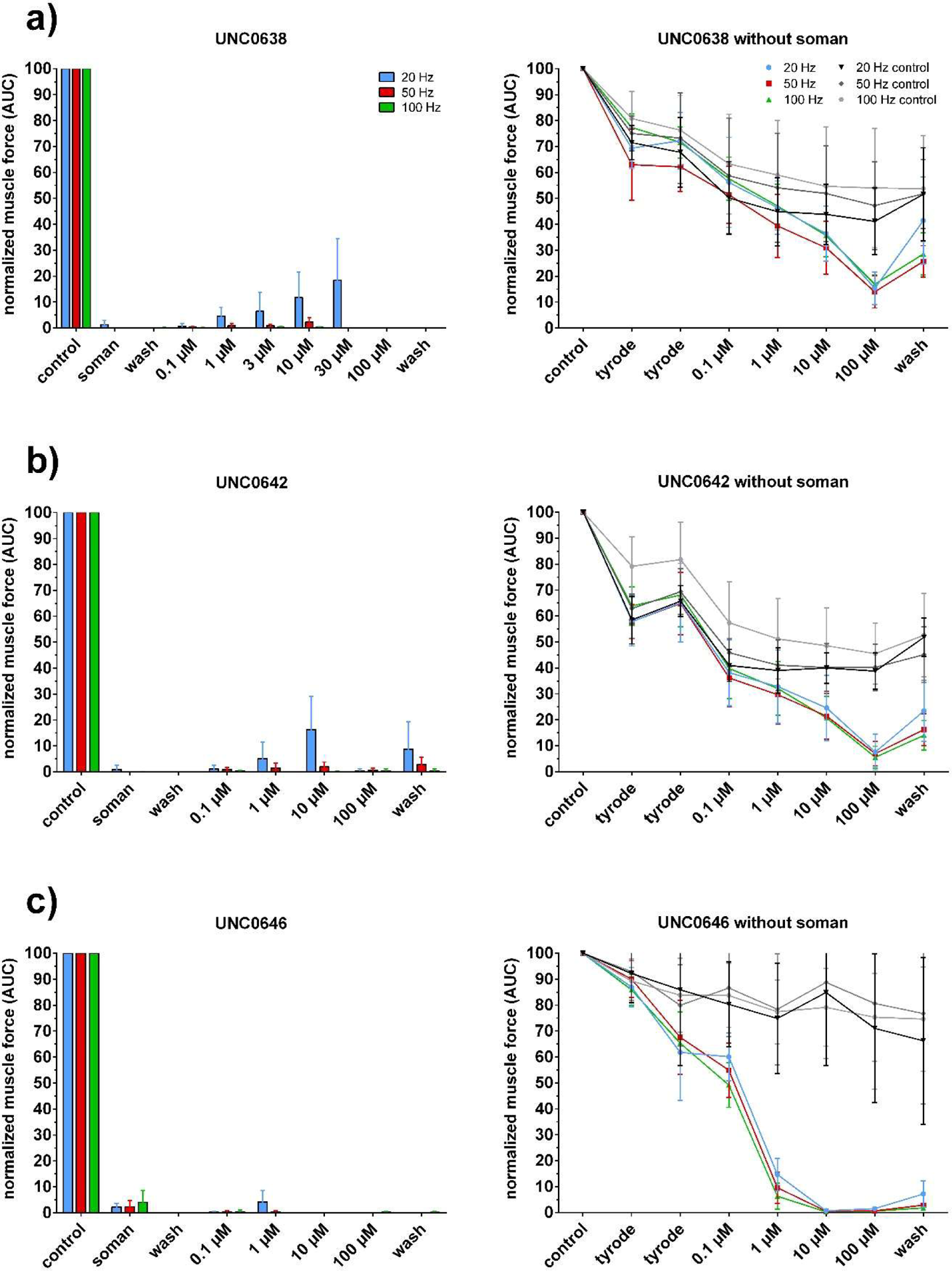
Muscle force of soman-poisoned (left) and un-poisoned (right) rat diaphragms after treatment with a) UNC0638, b) UNC0642, and c) UNC0646. Left: Muscle force of diaphragm muscle was blocked by 3 µM soman. Right: Muscle force generation of un-poisoned muscle. Muscle force generation was measured as the area under the curve normalized to muscle force under control conditions at the start of the measurement (n = 6 - 12).

Here, the muscle force of the functionally active muscle decreased when UNC0638 and UNC0642 were applied at high concentrations (i.e. ≥ 10 µM). This effect was even more pronounced for UNC0646, as it started at distinctly lower concentrations and led to a nearly complete inhibition of the muscle force at 1 µM (see Figure 6). Hence, the “bell-shape” of the curve of muscle-force recovery in soman-poisoned rat diaphragms upon treatment with UNC0638 and UNC0642 may be attributed to the above-described counteracting effect, which is, however, moderate, so that at lower concentration a positive effect still prevails. In contrast, in the case of UNC0646 no positive effect on muscle force recovery remains, as here the counteracting effect starts at distinctly lower concentrations and is more pronounced.

Finally, it should be mentioned, that the muscle force decreasing potential of the studied compounds appears to be reversible, as the muscle force partly recovers when the respective samples are subjected to a washing step (see Figure 6).

Overall, the quinazoline derivatives UNC0638, UNC0642 and UNC0646 identified as binders of the MB327-PAM-1 binding site of the nAChR are in principle also capable to restore muscle function of soman-poisoned muscle tissue. Though the maximum amplitudes for muscle force recovery of UNC0638 and UNC0642 are lower than those found for MB327, the maximum values are, remarkably, reached for the two quinazoline derivatives at distinctly lower concentrations than for MB327, which is likely to result from their higher binding affinities. Future studies will have to aim on a better understanding of the factors responsible for the counteracting effects regarding the recovery of muscle force in soman-poisoned rat diaphragms that might finally lead to more potent compounds.

## 3. Conclusion

The quinazoline-based compounds UNC0638, UNC0642, and UNC0646 had been identified as hits in a recent library screening campaign using competitive [^2^H_6_]MB327 MS Binding Assays in the search for new ligands addressing the MB327-PAM-1 binding site of the nAChR. In the present study, these compounds, which exhibit the highest affinities known so far for the MB327-PAM-1 binding site, have been used for the development of new MS Binding Assays for the aforementioned binding site of the nAChR with UNC0642 serving as reporter ligand, UNC0638 as internal standard and UNC0646 as competitor for the determination of non-specific binding. The new UNC0642 MS Binding Assays comprised the characterization of the binding of UNC0642 to the MB327-PAM-1 binding site of *Torpedo*-nAChR in saturation experiments and the determination of the binding affinity of a set of ligands of the aforementioned binding site in competition experiments. The results were in good accord with those obtained from the [^2^H_6_]MB327 MS Binding Assays, that had been used so far for the determination of binding affinities for the MB327 binding site. Carbachol used as a control had only a very small effect on reporter ligand binding in respective competition experiments. As this compound, carbachol, represents a ligand of the orthosteric binding site of the nAChR the observed effect on reporter ligand binding is likely to be attributed to an allosteric interaction between both binding sites. Based on the results obtained with the new UNC0642 MS Binding Assay it is reasonable to conclude, that this binding assay addresses the same binding site as the [^2^H_6_]MB327 MS Binding Assay, i.e. the MB327-PAM-1 binding site of the nAChR. Hence, the UNC0642 MS Binding Assays represent a valuable alternative to the [^2^H_6_]MB327 MS Binding Assays and profit from the high affinity of the reporter ligand, which will contribute to the robustness of the binding assays, and from the commercial availability of said compound. By using docking approaches and molecular dynamics simulations, the binding mode and key interactions of UNC0646 in the recently proposed allosteric binding site of the nAChR, the MB327-PAM-1 binding site, could be successfully unveiled.

*Ex vivo* studies revealed a beneficial effect of UNC0638 and UNC0642 on muscle force recovery of soman-poisoned rat diaphragms. In experiments with un-poisoned muscle tissues, a muscle force-reducing effect was uncovered for all three test compounds, which in the case of UNC0646 is likely to be so strong that it overcompensates the positive effect, that might originate from this compound on soman-poisoned muscle tissue, as well. This may explain the observed lack of such an effect for this compound. Though the maximum effect in muscle force recovery mediated by UNC0638 and UNC0642 was lower than that observed for MB327 and some analogs, this effect was reached at distinctly lower concentrations for UNC0638 and UNC0642 than that needed in the case of MB327. This might reflect the distinctly higher binding affinities of UNC0638 and UNC0642 for the MB327-PAM-1 binding site as compared to MB327 and analogs.

Overall, UNC0638, UNC0642, and UNC0646 have been found to address the MB327-PAM-1 binding site of the nAChR with high affinity, which renders quinazoline derivatives a promising class of compounds for further studies aiming at the development of drugs for the treatment of the nAChR-mediated pathological effects of organophosphorus poisoning.

## Experimental Section

### Materials

UNC0638 and UNC0642 (purity for both ≥ 95%) were purchased from MedChemExpress (Sollentuna, Sweden). UNC0646 (purity ≥ 99%) was received from Axon Medchem (Groningen, Netherlands) and carbachol (carbamoylcholine chloride, purity ≥ 98%) from Sigma Aldrich. MB327 and PTM0022 were synthesized in-house by Rappenglück et al., purities ≥ 95% (Rappenglück et al., 2018b). Frozen tissue of *Torpedo californica* electroplaque was purchased from Aquatic Research Consultants (San Pedro, CA, USA). Water was obtained from a Sartorius arium pro ultrapure water system (Sartorius, Göttingen, Germany) for all purposes. Organic solvents for LC-MS were received from VWR Prolabo (Darmstadt, Germany) in LS-MS grade. Ammonium formate as additive for LC-MS (purity ≥ 99%) was purchased from Sigma Aldrich (Taufkirchen, Germany). All other chemicals were purchased in analytical grade. Polypropylene reaction tubes and 96-deep well plates as well as pipette tips were received from Sarstedt (Nümbrecht, Germany).

### Preparation of nAChR-enriched membrane fragments

The nAChR-enriched membrane fragments were prepared from frozen electroplaque of *Torpedo californica* as described by Sichler et al. (Sichler et al., 2018).

### UNC0642 centrifugation-based MS Binding Assays

In general, for MS binding experiments with UNC0642 at *Torpedo*-nAChR, the reporter ligand was incubated with aliquots of the membrane preparation from *Torpedo californica* electroplaque (approx. 75 µg protein per sample) in incubation buffer (120 mM NaCl, 5 mM KCl, 8.05 mM Na_2_HPO_4_ and 1.95 mM NaH_2_PO_4_, pH 7.4). With a total volume of 1.25 mL, each binding sample was generated in a 1.5 mL reaction tube. Incubation took place in a shaking water bath (2 h, 25 °C). After that, the reaction tubes were centrifuged for 5 min at 4 °C and 23000 rpm (approx. 49000 x *g*, Heraeus Biofuge Stratos, rotor 3331, Thermo Scientific, Waltham, USA). In the next step, the formed pellets were freed from the supernatant using a Pasteur pipette, which was connected to a vacuum pump via a vacuum filter flask. Thereafter, pellets were washed two times by the addition of 1.5 mL ice-cold incubation buffer and instant removal of the latter by a vacuum-coupled Pasteur pipette. To liberate the bound reporter ligand, 500 µL acetonitrile (containing 500 nM UNC0638 as internal standard) were given to the pellets. The mixtures were subsequently subjected to ultrasound in an ultrasonic bath (SONOREX RK100, Bandelin electronic, Berlin, Germany) for 1 h. Next, the samples were vortexed intensively before they were centrifuged again under the same conditions as described before. Of the resulting supernatants 10 µL were transferred into a 96-deep well plate and diluted by the addition of 490 µL acetonitrile to each well. The 96-deep well plate was sealed with aluminum foil before the samples were finally analyzed via LC-ESI-MS/MS.

For saturation experiments, total binding was determined for fifteen reporter ligand concentrations, reaching from 200 nM to 100 µM. For the evaluation of non-specific binding, binding samples with the five lowest reporter ligand concentrations (200 nM - 1 µM) were, in addition, provided with an excess of competitor UNC0646 (100 µM). Regarding the amount of DMSO introduced by the stock solutions of the used compounds (10 mM in DMSO), all binding samples were adjusted to the same amount of 1% DMSO (*v*/*v*).

Competition experiments were performed in analogy to saturation experiments, except that the reporter ligand concentration in the binding samples was set to a fixed concentration of 1 µM. Furthermore, binding samples contained test compounds in general in six but at least in five different concentrations (100 nM - 10 mM). Total binding in the absence of any test compound was determined by means of control samples with only 1 µM of the reporter ligand UNC0642 present in addition to the *Torpedo* membrane preparation. For the determination of non-specific binding of the reporter ligand, binding samples containing 1 µM UNC0642 and *Torpedo* membrane preparation were additionally provided with 100 µM of UNC0646 as a competitor.

### Data Analysis

The concentration of the reporter ligand, UNC0642, in each sample was calculated by the Analyst software v. 1.6.1 (AB Sciex, Darmstadt, Germany) based on an underlying calibration curve. To ensure a reliable quantification, calibration standards, and a corresponding calibration function were generated for each binding experiment (see “Validation of the LC-ESI-MS/MS method” for details). Further analysis (e.g., linear regression, non-linear regression, and normalization) of the data to evaluate the binding experiments was done with the Prism software v. 6.07 (GraphPad software, La Jolla, CA, USA).

For saturation experiments, non-specific binding was determined only for the lowest five concentration levels (200 nM - 1 µM). By analyzing this data via linear regression forced through zero a linear regression function was established. This was subsequently used for the calculation of non-specific binding values for all reporter ligand concentrations applied in the assay. Subtracting non-specific binding from total binding yielded specific binding, which was further analyzed by the “One site binding (hyperbola)” regression tool to obtain the values for *B*_max_ and *K*_d_.

For competition experiments, the data received was firstly normalized with the total binding of the reporter ligand in the absence of a test compound being set to 100% and non-specific binding to 0%. The data was then analyzed with the “One site – fit Ki” regression tool, fixing top and bottom levels to 100% and 0% respectively, yielding competition curves. The derived IC_50_ values were automatically transformed into *K*_i_ values according to the Cheng-Prusoff equation by the additional input of the *K*_d_ value for the reporter ligand, UNC0642 (*K*_d_ = 6.7 µM), which was determined in saturation experiments as described above. If not stated otherwise, the results of binding experiments (*B*_max_, *K*_d_, *K*_i_) are given as means from three experiments ± SEM.

### LC-MS instrumentation

For preliminary experiments and method development, an API3200 triple quadrupole mass spectrometer with a TurboV-ESI source (Sciex, Darmstadt, Germany) was used in positive mode. After UNC0642 had been selected as a reporter ligand, all subsequent experiments were performed on an API5000. The two mass selectors, Q1 and Q3, were operated under unit resolution. For LC-ESI-MS/MS measurements the MS instrument was provided with an Agilent 1200 Series HPLC system (vacuum degasser G1379B, binary pump G1312B, oven G1316B, Agilent, Waldbronn, Germany). The stationary phase consisted of a YMC-Triart Diol-HILIC (50 mm x 2.0 mm, 3 µm; YMC Europe GmbH, Dinslaken, Germany) protected by two in-line filters (0.5 µm and 0.2 µm, IDEX, Wertheim-Mondfeld, Germany). The mobile phase consisted of a mixture of acetonitrile and an ammonium formate buffer (20 mM, pH 3.0) in a ratio of 80:20 (*v*/*v*). The flow rate amounted to 800 µL/min and the temperature of the column oven was set to 20 °C. For sample injection (injection volume: 10 µL), a HTS-PAL autosampler (CTC-Analytics, Zwingen, Switzerland) equipped with a 50 µL syringe was used. For the direct infusion of compound solutions into the ESI source, the HPLC system of the LC-ESI-MS/MS unit was exchanged by a syringe pump (Harvard Apparatus, Holliston, MA, USA).

### Establishing compound- and source-specific parameters

Mass transitions and compound-specific parameters for UNC0638, UNC0642, and UNC0646 were determined automatically by the “Compound optimization” tool of the Analyst software according to the manual of the respective mass spectrometer. For these experiments, the analytes were dissolved in a mixture of methanol and 0.1% aqueous formic acid (50:50 (*v*/*v*)). This solution was then directly introduced into the ESI source using a syringe pump. The optimized compound-dependent parameters for the determined mass transitions that were used for the detection of the analytes via MS/MS are listed in Table 3.

**Table 3.** Compound-specific parameters and corresponding mass transitions used for the detection of UNC0638, UNC0642, and UNC0646. DP = declustering potential, EP = entrance potential, CE = collision energy, CXP = cell exit potential.

| analyte | parent ion [M+H]<sup>+</sup><br>*m/z* | fragment ion<br>*m/z* | DP<br>[V] | EP<br>[V] | CE<br>[V] | CXP<br>[V] |
| --- | --- | --- | --- | --- | --- | --- |
| UNC0638 | 510.3 | 112.2 | 86 | 10 | 37 | 18 |
| UNC0642 | 547.3 | 112.1 | 80 | 10 | 43 | 18 |
| UNC0646 | 622.5 | 126.1 | 91 | 10 | 55 | 4 |

Source-specific parameters were optimized for the reporter ligand UNC0642 using the “Flow Injection Analysis” tool of the Analyst software, to which end a solution of acetonitrile containing 5 nM UNC0642 and 1:50 (*v*/*v*) matrix blank was repeatedly injected. The obtained parameters are as follows: collision gas (N_2_) = 6 psi, curtain gas (N_2_) = 20 psi, nebulizing gas (N_2_) = 30 psi, auxiliary gas (N_2_) = 50 psi, ion-spray voltage = 1500 V and temperature = 600 °C.

### Validation of the LC-ESI-MS/MS method

Matrix zero calibrator samples, which were necessary for the generation of calibration standards and quality control samples, were prepared in the same way as binding samples (see above: ‘UNC0642 centrifugation-based MS Binding Assays’) with the exception that the incubation was carried out in the absence of any compounds. Instead, matrix zero calibrator samples were later spiked with a defined amount of the reporter ligand, UNC0642, in order to generate the corresponding calibration standards and quality control samples. Thus, the dilution step prior to LC-ESI-MS/MS measurement was adapted (compared to binding samples) and 10 µL supernatant (containing no UNC0642, but like binding samples 500 nM of the internal standard, UNC0638) were diluted with 440 µL acetonitrile and another 50 µL of acetonitrile, which then contained an according amount of the reporter ligand, UNC0642. Following this procedure, quality control samples investigated the subsequently given concentration levels for the reporter ligand, UNC0642, with each prepared in six replicates: 50 pM (LLOQ), 500 pM, 5 nM, 50 nM. Calibration standards were studied in eight different reporter ligand concentration levels, each generated in three replicates (50 pM, 150 pM, 400 pM, 1.2 nM, 3.5 nM, 10 nM, 30 nM, 75 nM). The data for calibration standards was plotted in a coordinate system with peak area ratios of analyte vs. internal standard on the y-axis and the concentration ratios of analyte vs. internal standard on the x-axis. Calibration curves were then obtained by linear regression with a weighting factor of 1/x^2^.

Matrix blanks were prepared analogously to binding samples (see above: ‘UNC0642 centrifugation-based MS Binding Assays’) except that there were no compounds present during the incubation and with the difference, that acetonitrile without internal standard was added to the pellet after the washing process of the samples.

### Docking of UNC0646

The structure of the Torpedo nAChR (PDB-ID: 6UWZ (Rahman et al., 2020)) was used for docking. The α-neurotoxin and molecules from the crystallization buffer were removed, and the receptor was protonated using Protonate3D, as implemented in MOE v2020.09 (Chemical Computing Group, 2020) at pH 7. The termini were capped with *N*-methyl amide (NME) and acetyl (ACE) groups, respectively, using Maestro (Release 2022-3) (Schrödinger, 2021). The 3D conformation of the ligand was retrieved from the SMILES code and subsequently docked using MOE v2020.09 with default parameters for flexible docking (Chemical Computing Group, 2020).

### Molecular dynamics simulations

The nAChR in complex with two UNC0646 ligands, both at the negative (i.e., between the β- and α-subunit and the γ- and α-subunit) site of the α-subunit, was embedded in a membrane consisting of 1-palmitoyl-2-oleoyl-*sn*-glycero-3-phosphocholine (POPC) lipids and solvated in a rectangular box of “optimal point charge” (OPC) water using Packmol-Memgen (Schott- Verdaugo and Gohlke, 2019) from AmberTools22 (Case et al., 2023; Case et al., 2022). The edge of the box was set to be at least 12 Å away from the receptor atoms. KCl was added at a concentration of 150 mM and Cl- ions were used to neutralize the system. The AMBER22 package of molecular simulations software (Case et al., 2005) was used to perform MD simulations in combination with the ff19SB force field (Tian et al., 2020) for the protein and the Lipid21 force field (Dickson et al., 2022) for lipids. Ligand charges were calculated according to the RESP procedure (Bayly et al., 1993) with default parameters as implemented in antechamber (Wang et al., 2006) using electrostatic potentials generated with Gaussian16 (M. J. Frisch et al., 2016) at the HF 6-31G* level of theory; force field parameters for the ligand were taken from the gaff force field (Wang et al., 2004). Simulations were subsequently performed as described earlier (Kaiser et al., 2023). In short, first, a combination of steepest descent and conjugate gradient minimization was performed while lowering the positional harmonic restraints on receptor and ligand atoms from a force constant of 25 kcal mol^-1^ Å^-2^ to one of zero. Then, the system was stepwise heated to 300 K and, subsequently, positional harmonic restraints were decreased from a force constant of 25 kcal mol^-1^ Å^-2^ to one of zero.

Thereafter, 12 replicas of 500 ns length each of unbiased MD simulations were performed, using Langevin dynamics with a collision frequency of 2 ps^-1^ for temperature control and the Berendsen barostat with semi-isotropic pressure adaption. The trajectories were analyzed with CPPTRAJ (Roe and Cheatham, 2013). The per-residue effective binding energy was computed using the MM-PBSA method, as implemented in AMBER21 (Miller et al., 2012), in the presence of a heterogenous-dielectric implicit membrane model with spline fitting (Greene et al., 2019), an ionic strength of 0.15 M, and an internal dielectric constant of 4.

### Binding mode of UNC0642

Based on the MD simulations with UNC0646 bound to nAChR, we clustered the binding of UNC0646 to obtain a representative binding mode using the *k-means* algorithm, as implemented in CPPTRAJ (Roe and Cheatham, 2013). Based on the biggest cluster (containing 37% of all frames), we replaced the substituents of the quinazoline ring of UNC0646 according to the substitution pattern of UNC0642 and subsequently minimized the ligand in the presence of the receptor with all receptor atoms constrained using MOE v.2022.02 (Chemical Computing Group, 2023).

### Image generation

Images of nAChR were generated using PyMol v2.4.0 (Schrodinger, 2015).

### Alignment of subunits

Sequences of subunits were retrieved from the UniProt database (accessed on the 30^th^ of January, 2023) (The UniProt, 2023) and aligned using Jalview v2.11.2.6 (Waterhouse et al., 2009).

### Rat diaphragm myography

All procedures using animals followed animal care regulations. Preparation of rat diaphragm hemispheres from male Wistar rats (300 ± 50 g) and experimental protocol of myography was performed as described before with slight modifications (Seeger et al., 2012; Seeger et al., 2007). The stimulation was shortened from 25 Hz to 20 Hz and the pulsewidth from 50 to 10 µs. In short, for all procedures (including wash-out steps, preparation of soman and test compound solutions) aerated Tyrode solution (125 mM NaCl, 24 mM NaHCO_3_, 5.4 mM KCl, 1 mM MgCl_2_, 1.8 mM CaCl_2_, 10 mM glucose, 95% O_2_, 5% CO_2_; pH 7.4; 25 ± 0.5 °C) was used. After the recording of control muscle force one hour after preparation, the muscles were incubated in the Tyrode solution, containing 3 μM soman for 20 min. Following a 20 min wash-out period, the test compounds were added in ascending concentrations (0.1 μM to 100 μM). The incubation time was 20 min for each concentration. The electric field stimulation was performed with 10 μs pulse width and 2 A amplitudes. The tetanic stimulation of 20 Hz, 50 Hz, 100 Hz were applied for 1 s and in 10 min intervals. Muscle force was calculated as a time-force integral (area under the curve, AUC) and constrained to values obtained for maximal force generation (muscle force in the presence of Tyrode solution without any additives; 100%). All results were expressed in means ± SD (n = 6 - 12). For all data analysis, Prism 5.0 (GraphPad Software, San Diego, CA, USA) was used.

## Acknowledgments

This work was supported by the German Ministry of Defence (E/U2AD/KA019/IF558). We are grateful for computational support and infrastructure provided by the “Zentrum für Informations- und Medientechnologie” (ZIM) at the Heinrich Heine University Düsseldorf and the computing time provided by the John von Neumann Institute for Computing (NIC) to HG on the supercomputer JUWELS at Jülich Supercomputing Center (JSC) (user IDs: HKF7, VSK33, nAChR).

