## Supplementary material for "MS Binding Assays with UNC0642 as reporter ligand for the MB327 binding site of the nicotinic acetylcholine receptor": SI

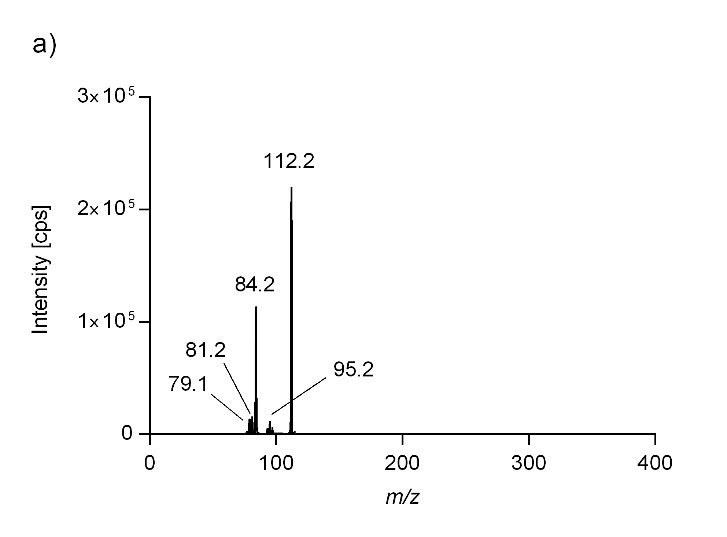

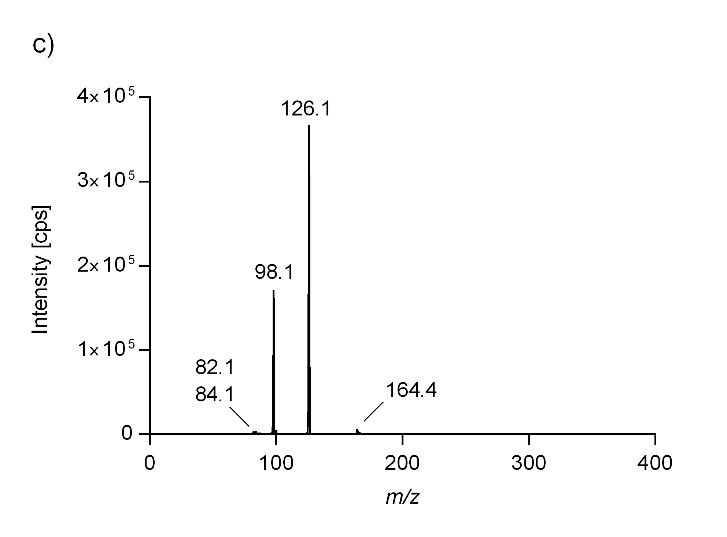

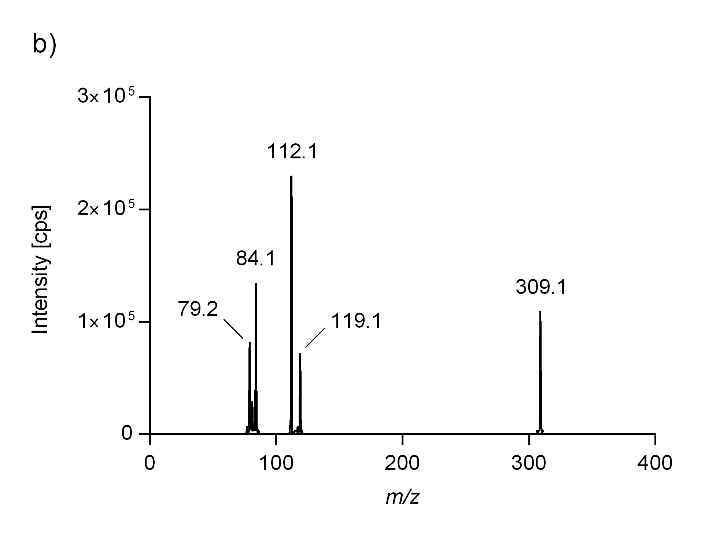
**Supporting Information**

**Figure S1.** Product ion spectra of UNC0638 (a), UNC0642 (b), and UNC0646 (c).

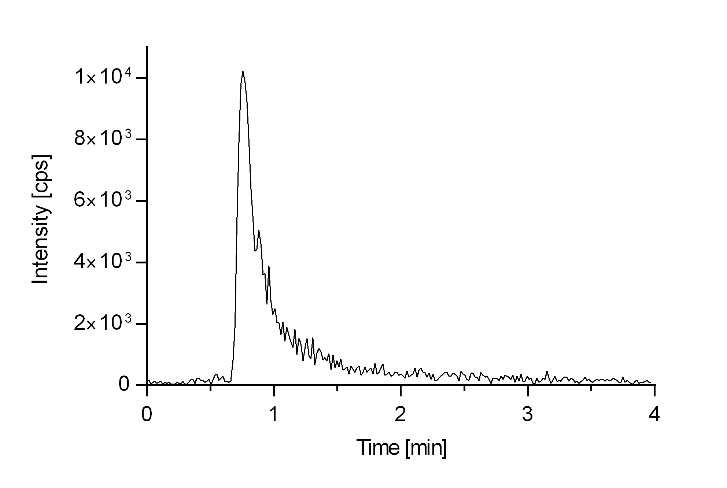
**Figure S2.** Representative LC-ESI-MS/MS-MRM chromatogram of UNC0646 (*m*/*z* 622.5/126.1, 10 nM in acetonitrile) recorded at an API3200, using the following chromatographic parameters. Stationary phase: YMC-Triart Diol-HILIC (50 mm x 2.0 mm, 3 µm). Mobile Phase: 65:35 (*v*/*v*) acetonitrile and ammonium formate buffer (20 mM, pH 3.0), 20 °C. Flow rate: 600 µL/min. Injection volume: 10 µL.

**Table S1.** Validation results of the LC-ESI-MS/MS method for the quantification of UNC0642

| **Sample (n)** | **Intra-Series** | | | | | | | | | **Inter-Series** | | |
| --- | --- | --- | --- | --- | --- | --- | --- | --- | --- | --- | --- | --- |
|  | **Series 1** | | | **Series 2** | | | **Series 3** | | |  |  |  |
|  | **M** | **A** | **P** | **M** | **A** | **P** | **M** | **A** | **P** | **M** | **A** | **P** |
| 50 pM Cal (3) | 0.0502 | 100 | 11.4 | 0.0514 | 103 | 11.1 | 0.0507 | 101 | 9.51 | 0.0508 | 102 | 9.31 |
| 150 pM Cal (3) | 0.148 | 98.7 | 0.676 | 0.141 | 93.8 | 7.67 | 0.144 | 96.0 | 5.24 | 0.144 | 96.1 | 5.08 |
| 400 pM Cal (3) | 0.399 | 99.7 | 4.40 | 0.375 | 93.8 | 3.53 | 0.386 | 96.4 | 4.31 | 0.386 | 96.6 | 4.45 |
| 1.2 nM Cal (3) | 1.22 | 102 | 2.96 | 1.23 | 102 | 0.471 | 1.28 | 106 | 0.452 | 1.24 | 103 | 2.62 |
| 3.5 nM Cal (3) | 3.41 | 97.3 | 4.78 | 3.54 | 101 | 5.67 | 3.33 | 95.2 | 1.25 | 3.43 | 97.9 | 4.65 |
| 10 nM Cal (3) | 11.1 | 111 | 2.38 | 10.9 | 109 | 1.92 | 11.2 | 112 | 3.73 | 11.0 | 110 | 2.72 |
| 30 nM Cal (3) | 27.9 | 92.9 | 2.10 | 28.5 | 95.1 | 1.42 | 27.8 | 92.6 | 1.62 | 28.1 | 93.5 | 1.98 |
| 75 nM Cal (3) | 73.6 | 98.1 | 1.13 | 77.1 | 103 | 1.40 | 74.7 | 100 | 2.70 | 75.1 | 100 | 2.63 |
| Equation of the calibration curve ^a^ | y = 1.57 * x + 0.00292 (r = 0.9973) | | | y = 1.74 * x + 0.00217 (r = 0.9971) | | | y = 1.86 * x + 0.00182 (r = 0.9914) | | | - | | |
| 50 pM QC (6) | 0.0471 | 94.2 | 5.45 | 0.0467 | 93.5 | 6.51 | 0.0502 | 100 | 5.15 | 0.0480 | 96.0 | 6.31 |
| 500 pM QC (6) | 0.484 | 96.8 | 3.84 | 0.467 | 93.5 | 5.80 | 0.460 | 91.1 | 4.89 | 0.470 | 94.1 | 5.09 |
| 5 nM QC (6) | 4.93 | 98.6 | 2.60 | 5.16 | 103 | 5.28 | 5.03 | 101 | 2.46 | 5.04 | 101 | 4.01 |
| 50 nM QC (6) | 47.5 | 95.0 | 2.69 | 49.1 | 98.0 | 5.68 | 46.0 | 92.1 | 2.66 | 47.6 | 95.1 | 4.66 |

**M** = mean concentration [nM], **A** = accuracy [%], **P** = precision [%], ^a^ weighting factor: 1/x^2^

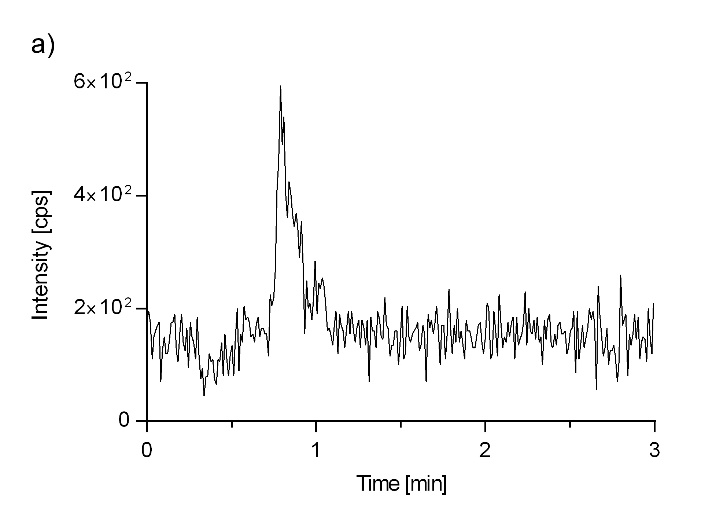

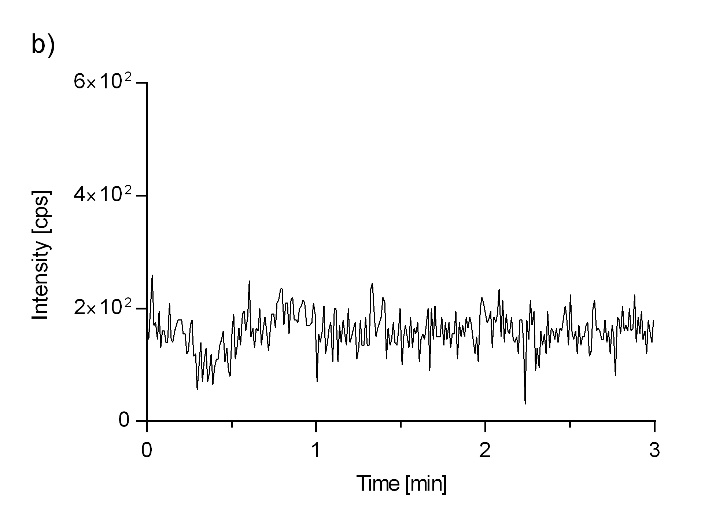
**Figure S3.** Representative LC-ESI-MS/MS-MRM chromatograms of UNC0642 (*m*/*z* 547.3/112.1) recorded as part of the validation process of the applied LC-ESI-MS/MS quantification method. Chromatographic conditions: stationary phase, YMC-Triart Diol-HILIC (50 mm x 2.0 mm, 3 µm); mobile phase, 80:20 (*v*/*v*) acetonitrile and ammonium formate buffer (20 mM, pH 3.0), 20 °C; flow rate: 800 µL/min; injection volume: 10 µL. (a) LLOQ of 50 pM, (b) Matrix blank.

**Figure S4.** Conformations of UNC0646 in MB327-PAM-1 in each subunit after blind docking experiments. a) Extracellular view of the receptor. The two α-subunits are colored yellow and green, the δ-subunit pink, the β-subunit cyan, and the γ-subunit salmon. Binding mode of UNC0646 in between the b) γ- and α-subunit, c) α- and δ-subunit, d) δ- and β-subunit, e) β- and α-subunit, and f) α- and γ-subunit. The receptor is shown as ribbon, the surface as a grey contour, and the ligand as sticks.

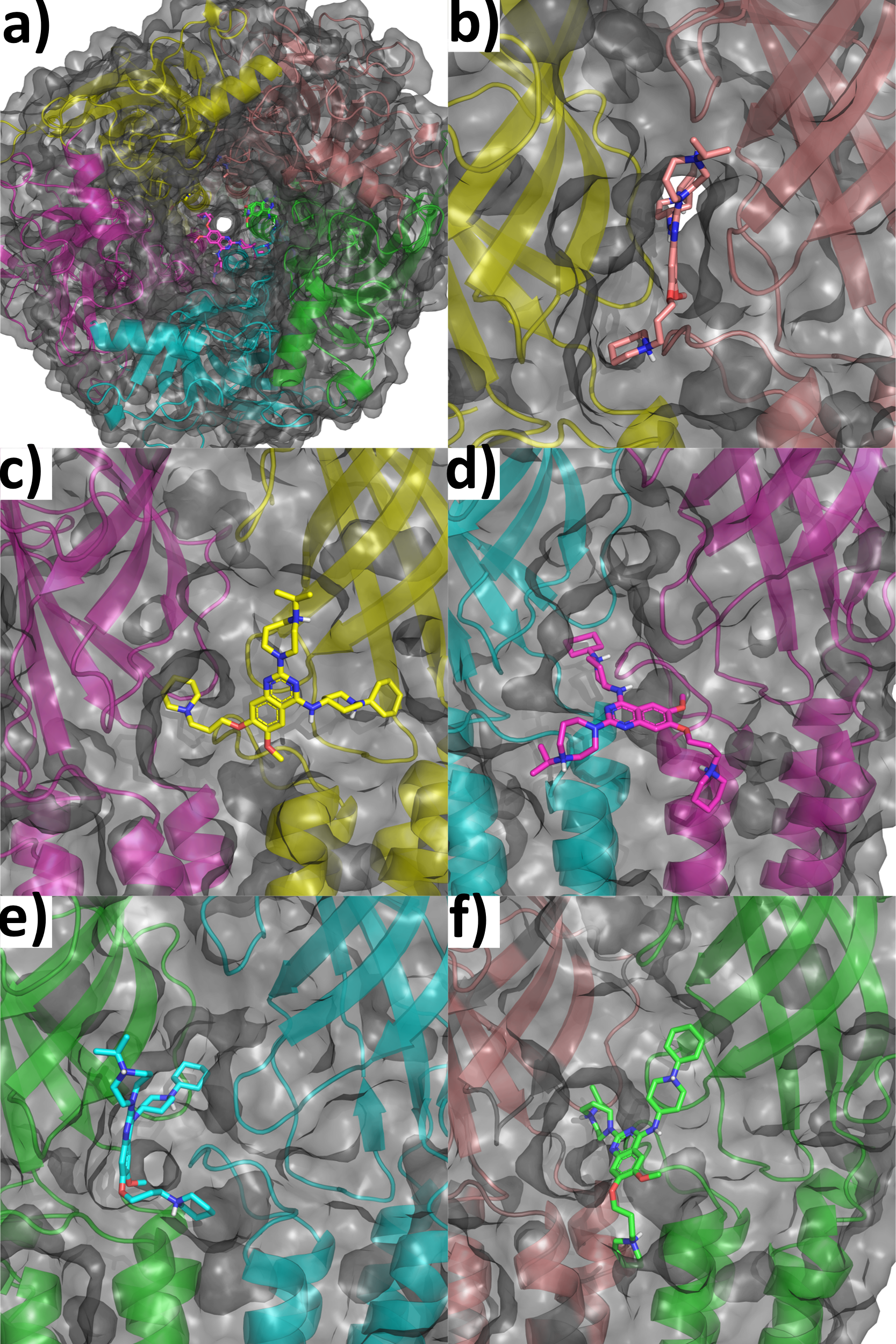

**Figure S5.** Docked orientation of UNC0646 between the γ- and α-(yellow) subunit. The γ-subunit has been omitted for clarity. The substituent in position 4 of the quinazoline ring is oriented in a) in a boat-twist conformation in the best-ranked pose and b) in a chair-chair conformation in the second-best-ranked conformation.

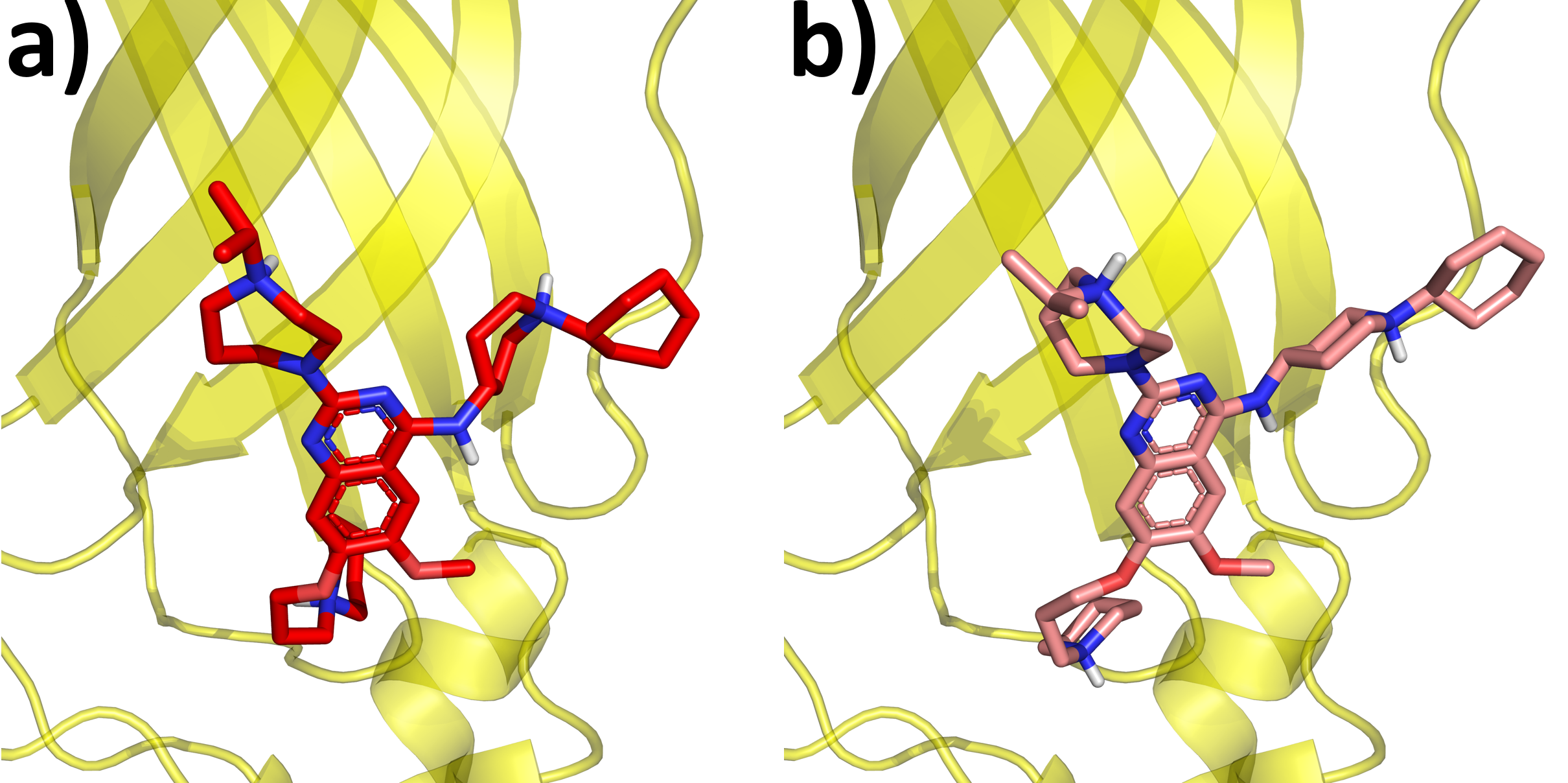

**Figure S6.** Docked binding mode of UNC0646 in between the β- and α-subunit. The ligand is shown in cyan and side chains similar to those in Figure 1 are shown in forest green.

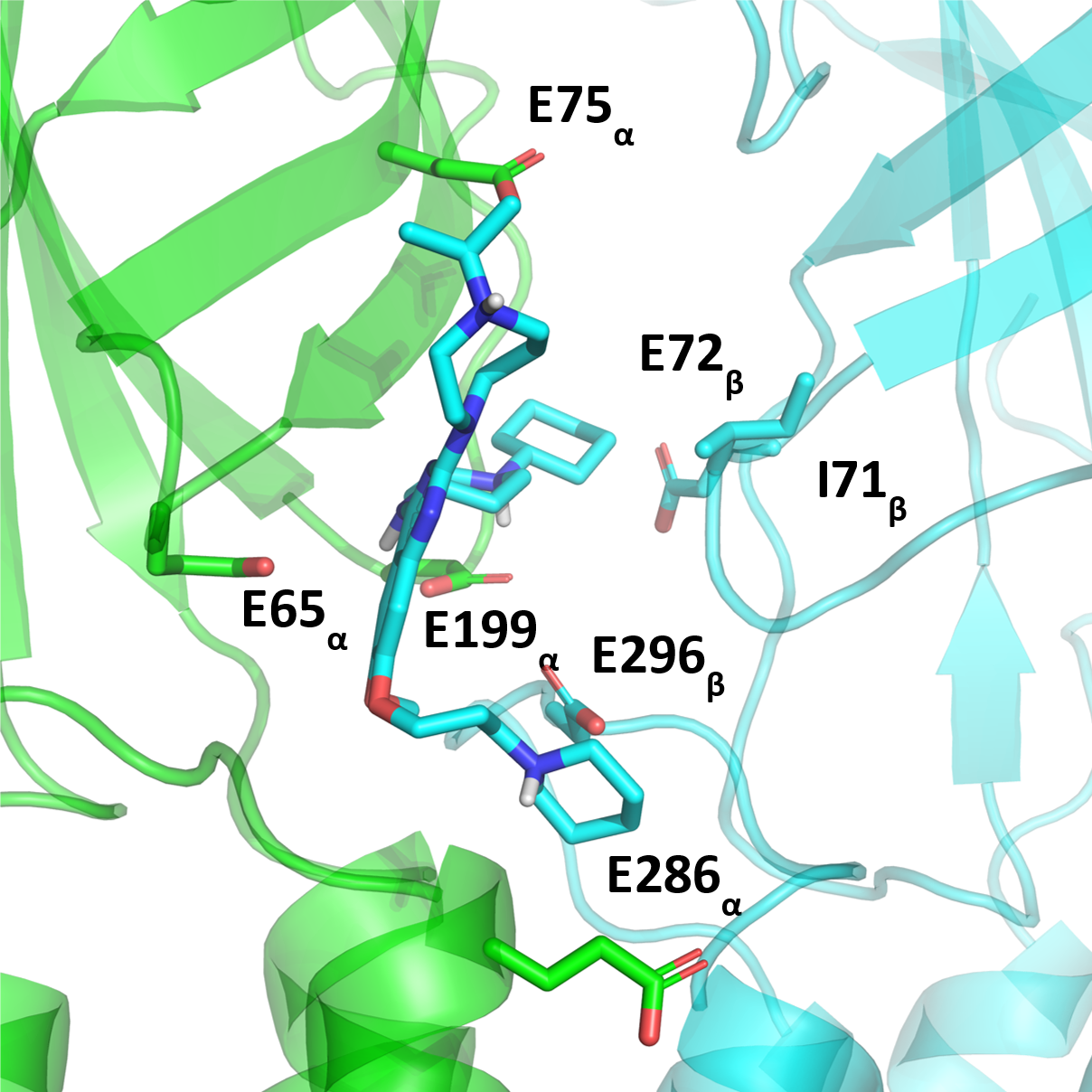

**Figure S7.** Backbone (C, CA, N) RMSD of nAChR during 12 replicas of 500 ns long MD simulations compared to the first frame of the production runs.

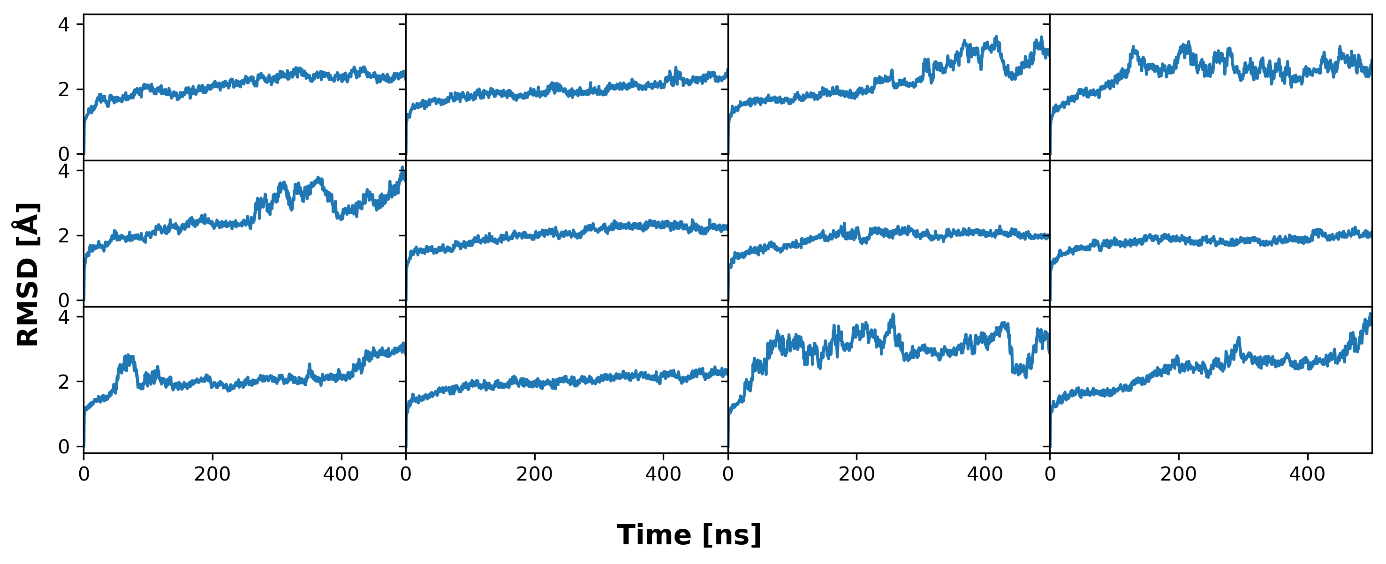

**Figure S8.** Electron density of water and membrane components over all 12 replicas of 500 ns long MD simulations. The electron density is plotted against the z-coordinate with the membrane centered at 0 Å for phosphatidylcholine (PC), oleic acid (OL), palmitoyl acid (PA), and water (WAT).

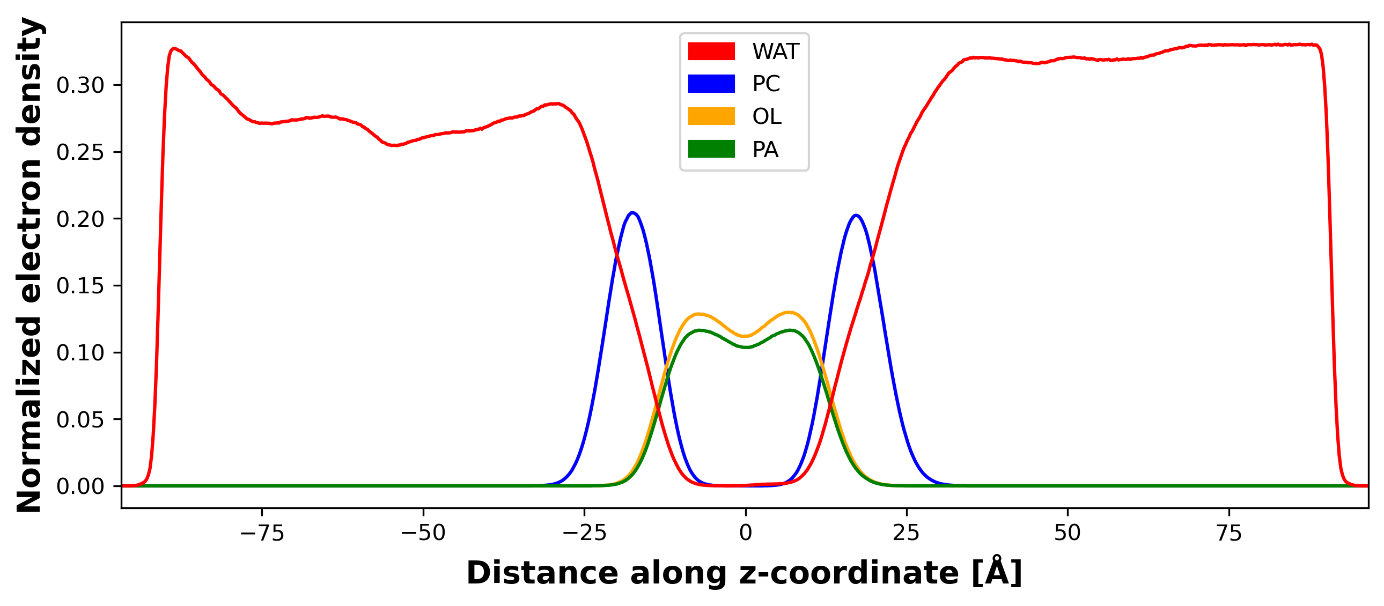

**Figure S9.** MMPBSA per-residue decomposition of the binding effective energy of UNC0646 in the subunits composing binding site A (γ- and α-subunit). Glutamates are shown in red.

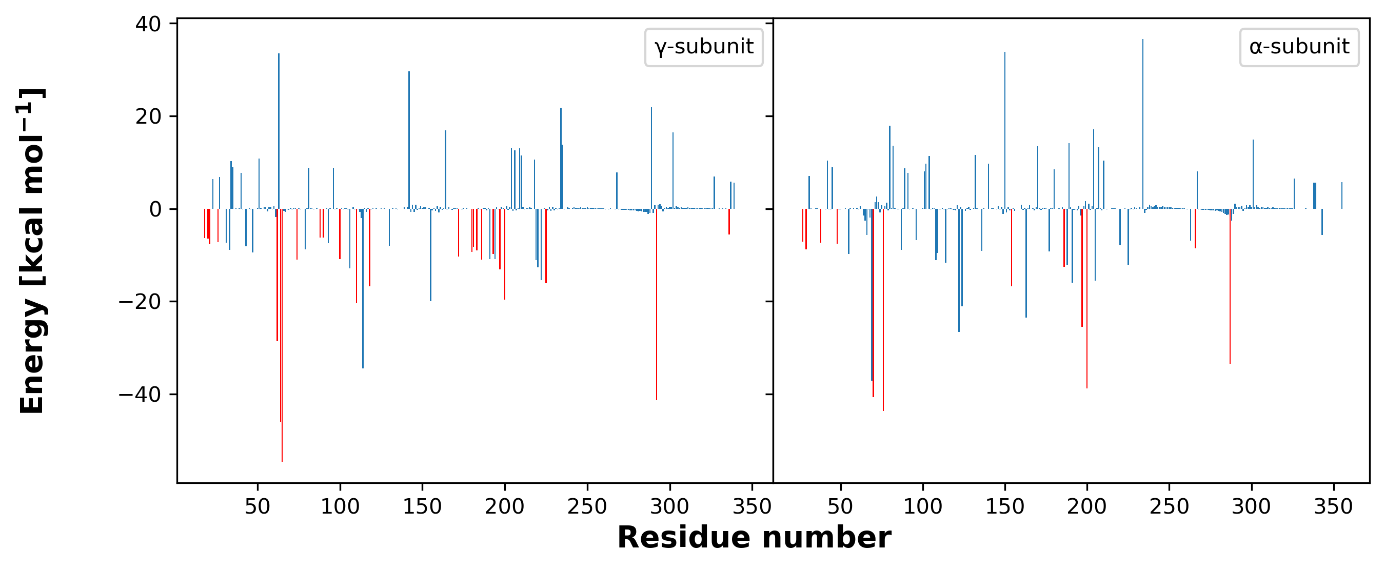

**Figure S10.** Binding mode of UNC0642 in MB327-PAM-1. The binding mode was obtained from a representative binding mode of UNC0646 in binding site A (between the γ- and α-subunit) and subsequent minimization of the ligand in the binding site.

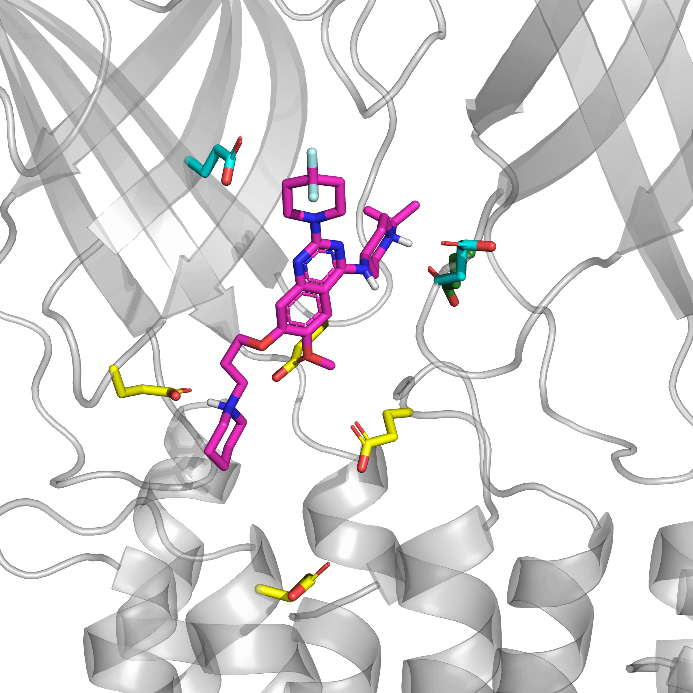
